# Boosting NADP-malic enzyme 1 enhances seed vigor and longevity in *Arabidopsis thaliana*

**DOI:** 10.64898/2026.04.16.718918

**Authors:** Ying Fu, Maroua Bouzid, Eva-Maria Schulze Isig, Maximilian Klamke, Gereon Poschmann, Martin Sosa, Mariel Gerrard Wheeler, Mariana Saigo, Veronica G. Maurino

**Affiliations:** Molecular Plant Physiology, Institute of Cellular Molecular Botany (IZMB), University of Bonn, Kirschallee 1, 53115 Bonn, Germany; Molecular Proteomics Laboratory, Biomedical Research Centre (BMFZ) & Institute of Molecular Medicine, Proteome research, Medical Faculty and University Hospital Düsseldorf, Heinrich Heine University Düsseldorf, Universitätsstraße 1, 40225 Düsseldorf, Germany; Centro de Estudios Fotosintéticos y Bioquímicos (CEFOBI-CONICET), University of Rosario, Suipacha 570, 2000 Rosario, Argentina

## Abstract

Seed longevity is a key determinant of crop establishment, productivity, and germplasm conservation. During storage and germination, reactive oxygen species accumulate and contribute to seed aging through oxidative damage and loss of viability. The maintenance of redox homeostasis therefore relies on NADPH-dependent antioxidant systems, which require a continuous supply of reducing power. NADP-dependent malic enzyme 1 (NADP-ME1), represents a source of NADPH supporting antioxidant defense during seed aging. Here, we show that enhanced expression of *NADP-ME1* positively contributes to seed vigor and longevity in *Arabidopsis thaliana*. *NADP-ME1* overexpression lines exhibited faster germination and higher overall germination after accelerated aging, whereas knockout mutants showed markedly reduced germination performance. Enhanced post-aging vigor in the overexpression lines was associated with reduced oxidative damage as indicated by lower malondialdehyde and hydrogen peroxide accumulation, along with preservation of specific polyunsaturated fatty acids, and increased γ-tocopherol levels in aged dry seeds. Enhanced expression of *NADP-ME1* reshapes the transcriptome of germinated seeds under fresh conditions compared with the wild type, while only minimal differences between genotypes are detected in aged seeds. These results suggest that NADP-ME1 contributes to the establishment of a transcriptional state associated with enhanced seed vigor and improved post-aging germination. Finally, co-immunoprecipitation coupled to mass spectrometry and bimolecular fluorescence complementation identified aspartate aminotransferase 2 as a NADP-ME1 interactor, pointing to a link between malate metabolism and amino acid-related metabolic adjustment. Together, these results identify NADP-ME1 as a determinant of seed resilience to aging and a potential target for improving seed quality.

## Introduction

Seeds are central to agriculture, serving as the primary means of crop propagation and as reservoirs of genetic diversity for future plant generations. They are the foundation of global food security, enabling the storage and dissemination of crops across time and space (Gupta et al., 2022). The quality and viability of seeds are therefore critical not only for individual farmers but also for the agricultural economy and food supply chains worldwide. However, despite careful handling and storage under controlled conditions, seeds inevitably lose their vigor and viability over time, a phenomenon influenced by both intrinsic genetic factors and external environmental conditions (Kurek et al., 2019; Rajjou and Debeaujon, 2008). This gradual decline in seed quality poses substantial challenges, such as, reduced germination rates, delayed seedling establishment, and compromised plant growth which can lead to significant yield losses, directly impacting economic returns and agricultural sustainability (Finch-Savage and Leubner-Metzger, 2006).

Seed deterioration, or aging, is a complex, multifactorial process that begins at the molecular level. During storage and early germination, seeds undergo progressive changes in cellular structures, enzyme activity, and metabolic pathways (Bailly et al., 1996; Zhang et al., 2021). Protective mechanisms, including seed coat integrity, heat shock proteins, late embryogenesis abundant (LEA) proteins, non-reducing sugars, flavonoids, and tocopherols, help stabilize cellular structures and mitigate damage; however, these defenses are gradually overwhelmed as aging advances (Rajjou and Debeaujon, 2008; Renard et al., 2020a; Sattler et al., 2004; Zinsmeister et al., 2020). One of the primary drivers of seed deterioration is the accumulation of reactive oxygen species (ROS). ROS inevitably accumulate during desiccation, storage, and germination (Chen et al., 2012; Wojtyla et al., 2016). The most common ROS in plants include singlet oxygen (¹O_2_), hydrogen peroxide (H_2_O_2_), superoxide anion (O_2_^•-^), and hydroxyl radical (OH•) (Bailly, 2004; Chen et al., 2019). During imbibition, the shift from quiescence to active metabolism is accompanied by increased respiration, which enhances electron leakage from the mitochondrial electron transport chain, generating superoxide anions that are subsequently converted to H_2_O_2_ by superoxide dismutase (Zhang et al., 2021). Excessive ROS induce lipid peroxidation, particularly of polyunsaturated fatty acids (PUFAs) in cellular membranes and storage lipids, as well as protein oxidation, enzyme inactivation, and DNA damage (Bailly, 2004; Lehner et al., 2008; Manisha Nigam et al., 2019; Pukacka and Ratajczak, 2007; Wojtyla et al., 2016). Consequently, aged seeds exhibit increased electrolyte leakage, elevated malondialdehyde (MDA) and lipid hydroperoxides, and decreased concentrations of antioxidant metabolites such as ascorbate and glutathione, as reported across multiple species under both natural and accelerated aging conditions (Goel and Sheoran, 2003; Naghisharifi et al., 2024; Ratajczak et al., 2015; Tonguç et al., 2023; Xia et al., 2015; Xin et al., 2014).

Disruption of ROS homeostasis accelerates seed deterioration, as evidenced by the reduced longevity of *Arabidopsis* mutants in dehydroascorbate reductase (DHA), whereas mutants with reduced NADPH oxidase (RBOHs) activity, and consequently lower ROS production, show enhanced longevity (Renard et al., 2020b). Conversely, improving oxidative stress tolerance enhances seed viability, as demonstrated in transgenic tobacco overexpressing Cu/Zn-superoxide dismutase and ascorbate peroxidase (Lee et al., 2010), as well as in mutants affecting lipid oxidation (OsLOX10; Wang et al., 2023b), hormone signaling (BG14; Wang et al., 2023a), and protein turnover (ATL5; He et al., 2023), highlighting the importance of coordinated mechanisms that limit oxidative damage and maintain cellular homeostasis. In addition to enzymatic antioxidant systems, non-enzymatic antioxidants play a crucial role in maintaining ROS homeostasis and protecting seeds from oxidative damage. Tocopherols and other tocochromanols, act as chain-breaking scavengers of lipid peroxyl radicals, and their deficiency severely reduces seed longevity (Nguyen et al., 2015; Rehmani et al., 2022; Sattler et al., 2004; Vom Dorp et al., 2015). Flavonoids in the seed coat also contribute to oxidative protection, as demonstrated by reduced germination and increased abnormalities in flavonoid-deficient testa mutants (Debeaujon et al., 2000; Rajjou and Debeaujon, 2008). Collectively, these findings support a model in which seed aging reflects a progressive imbalance between ROS production and detoxification, leading to irreversible oxidative damage, programmed cell death, and loss of viability.

NADPH is a major source of reducing equivalents for antioxidant systems and cell growth and is thus involved in diverse processes such as pathogen defense and root elongation (Corpas and Barroso, 2014; Foreman et al., 2003; Fu et al., 2026; Voll et al., 2012). NADP-dependent malic enzyme (NADP-ME) contributes to this process by catalyzing the oxidative decarboxylation of malate to pyruvate, thereby generating NADPH. In *Arabidopsis*, *NADP-ME1* (*AT2G19900*) is predominantly expressed in the cytosol of embryos and roots, with its expression later becoming restricted to secondary roots (Fu et al., 2026; Gerrard Wheeler et al., 2005). During seed maturation, NADP-ME1 is the main isoform contributing to total NADP-ME activity, and seeds lacking this isoform exhibit increased protein carbonylation, and reduced germination under both natural and accelerated aging conditions (Arias et al., 2018; Yazdanpanah et al., 2019).

Taken together, the available evidence points to NADP-ME1 as an important determinant of seed longevity. Yet, its precise contribution to seed survival during aging remains insufficiently defined. To address this gap, we examined wild-type, loss-of-function mutants, and overexpression lines under fresh and accelerated aging conditions. Through a combined physiological, biochemical, and transcriptomic approach, we aimed to determine how NADP-ME1 influences redox homeostasis and oxidative damage during seed aging, and to establish its importance in maintaining seed viability. Given the importance of seed longevity for storage, crop establishment, and agricultural productivity, NADP-ME1 stands out as a promising target for the development of strategies to enhance seed viability.

## Results

### Germination and seed viability assays in *NADP-ME1* knockout mutants and overexpression lines

To investigate the contribution of NADP-ME1 to seed viability and responses to aging, we generated four independent *NADP-ME1* overexpression lines (OX1-OX4) carrying the full-length *NADP-ME1* cDNA under the control of the CaMV35S promoter. All four lines exhibited a 4- to 6-fold increase in *NADP-ME1* transcript abundance in roots relative to the WT (Fig. 1A). By contrast, the T-DNA insertion lines (ko lines: *me1.1-me1.3*), generated in our previous study (Fu et al., 2026), showed significantly reduced *NADP-ME1* transcript levels in root compared with the WT. Based on these expression patterns, the overexpression lines OX2 and OX3, together with the ko lines *me1.2* and *me1.3*, were selected for further analyses.

**Figure 1.**
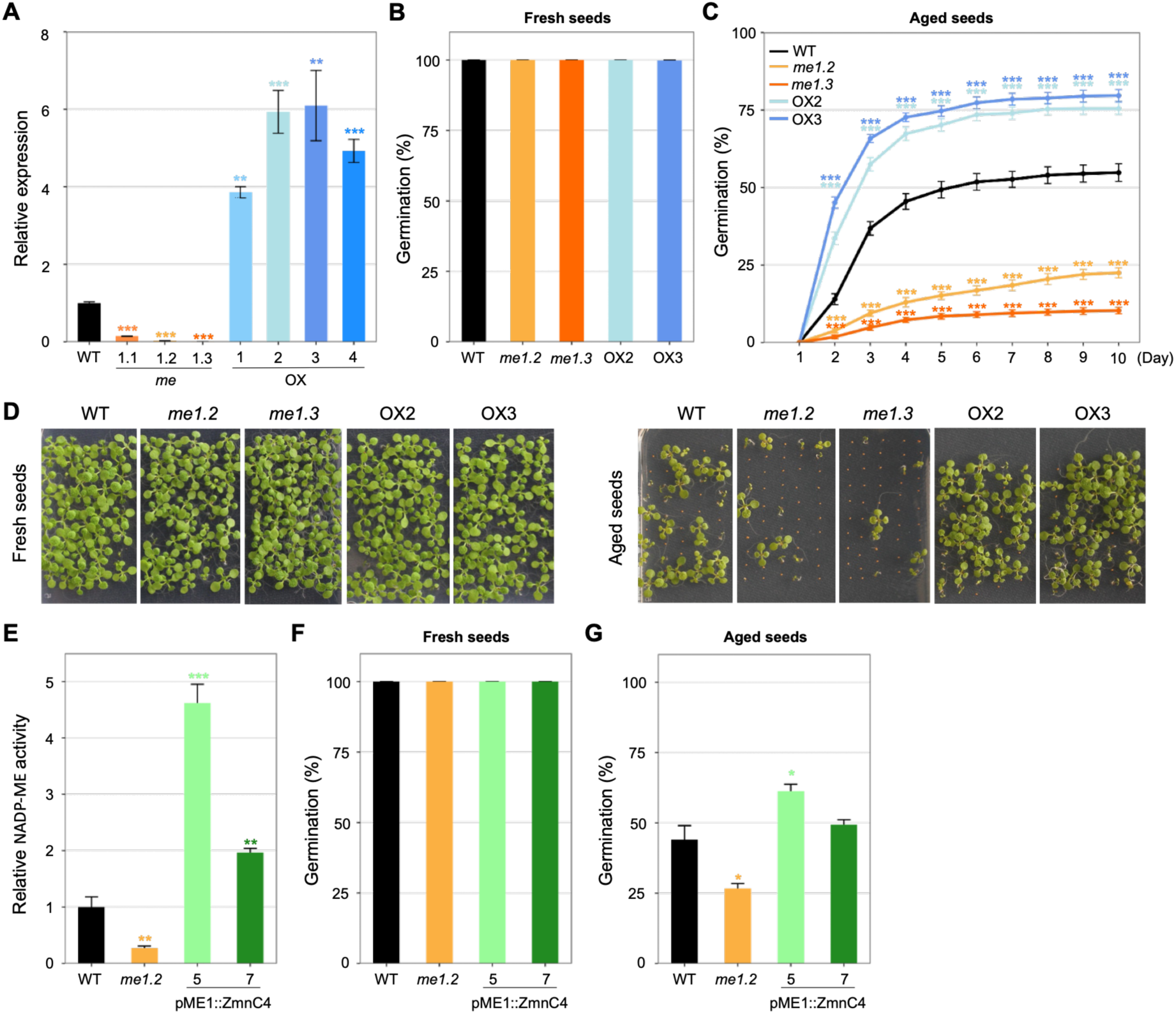
Transcript levels of *NADP-ME1* in different genotypes and seed viability assays in fresh and aged seeds. **A)** *NADP-ME1* transcript abundance in roots of WT, three independent knockout mutants (*me1.1*-*me1.3*), and four independent overexpression lines (OX1-OX4, driven by the CaMV35S promoter), quantified by qPCR. Transcript levels were normalized to *ACTIN2* (*AT3G18780*). **B)** Germination percentage of fresh seeds of WT, knockout mutants (*me1.2* and *me1.3*), and overexpression lines (OX2 and OX3) on ½ MS medium, assessed at 2 days after imbibition. Germination was assessed by counting seeds with visible radicles. **C)** Germination percentage of aged seeds from the same genotypes in B on ½ MS medium, monitored over multiple days after the imbibition phase. Data are presented as means ± SE (n = 4). **D)** Representative images of fresh or aged seeds from the same genotypes in B grown on ½ MS medium for 10 days. **E)** Relative NADP-ME activity in extracts of fresh dry seeds from WT, *me1.2*, and the transgenic lines pME1::ZmnC4-5 and pME1::ZmnC4-7. **F)** Germination percentage of fresh seeds from the same genotypes in E assessed at 2 days after imbibition. **G)** Germination percentage of aged seeds from the same genotypes as in E assessed at 10 days after imbibition. Data are presented as means ± SE (n = 3). Asterisks indicate significant differences compared to WT (* *p* < 0.05; ** *p* < 0.01; *** *p* < 0.001; Student’s t-test).

Fresh seeds of the WT, ko, and OX genotypes all reached 100% germination within 2 days after 48 h of imbibition and transfer to the growth chamber (Fig. 1B). However, marked differences emerged after accelerated aging treatment (Fig. 1C and D). WT seeds showed reduced vigor, reaching a maximum germination of approximately 53% by 7 days after imbibition. Germination was much more severely impaired in the ko lines, with *me1.3* reaching only 10% and *me1.2* approximately 19% at the same time point. By contrast, the OX lines displayed enhanced germination, attaining approximately 75% by day 7. Consistent with their enhanced vigor, seeds of the OX lines reached 50% germination within 2 days of transfer, whereas WT seeds reached only about 10% and seeds of the ko lines remained below 4% at the same time point.

To assess whether this phenotype could also be conferred by a different NADP-ME isoform, we generated two transgenic lines, pME1::ZmnC4-5 and pME1::ZmnC4-7, expressing in the cytosol the maize nonC4-NADP-ME (Alvarez et al., 2019; Saigo et al., 2004) under the control of the AtNADP-ME1 promoter. NADP-ME activity in fresh dry seeds, expressed relative to the WT level (WT = 1), increased 2.0-fold and 4.6-fold in pME1::ZmnC4-7 and pME1::ZmnC4-5, respectively (Fig. 1E), whereas it was reduced by approximately 73% in *me1.2*. Germination of fresh seeds was 100% in all genotypes (Fig. 1F). Following accelerated aging, germination reached 42% in WT, 27% in *me1.2*, 49% in pME1::ZmnC4-7, and 61% in pME1::ZmnC4-5, with the latter significantly exceeding the WT value (Fig. 1G).

These findings support a positive role for *NADP-ME1* in seed longevity and indicate that enhanced NADP-ME activity improves tolerance to seed aging.

### Altered seed vigor is not associated with changes in seed physical traits, soluble protein content, or glycation

To determine whether the differences in seed vigor observed after accelerated aging were associated with changes in basic seed physical properties or in general protein quality traits, we measured seed fresh weight, dry weight, water content, total soluble protein content, and Amadori products in fresh and aged seeds of the different genotypes. Seed fresh weight, dry weight, and water content did not differ significantly either between fresh and aged seeds or among genotypes (Suppl. Fig. 1A-C). Total soluble protein content decreased markedly upon imbibition compared with dry seeds, regardless of genotype or aging treatment (Suppl. Fig. 1D). Likewise, Amadori products increased during imbibition relative to dry seeds, but no significant differences were detected among genotypes or between fresh and aged seeds (Suppl. Fig. 1E). Together, these results indicate that changes in protein turnover and glycation are primarily associated with the developmental transition from dry to imbibed seeds, while the beneficial effect of NADP-ME1 on seed vigor is observed without compromising seed physiological quality.

### NADP-ME1 influences lipid peroxidation and ROS accumulation during germination of aged seeds

Lipid peroxidation and H_2_O_2_ accumulation were measured to assess oxidative damage associated with seed aging. In fresh seeds, neither malondialdehyde (MDA) nor H_2_O_2_ levels differed significantly among WT, ko, and OX lines at different stages (dry, imbibed, or 2 days after imbibition), indicating that NADP-ME1 expression does not affect basal oxidative status (Fig. 2A and C). After accelerated aging, oxidative stress increased markedly during germination, as shown by the higher MDA and H_2_O_2_ levels in aged seeds relative to fresh seeds (Fig. 2B and D). This increase was particularly pronounced in the ko mutants, which accumulated significantly more MDA and H_2_O_2_ during germination than WT and OX lines (Fig. 2A and C). During imbibition, lipid peroxidation also increased significantly in aged seeds of WT and ko lines, whereas the OX lines did not show this increase and maintained lower MDA levels than the mutants (Fig. 2B). Together, these results indicate that oxidative damage associated with seed aging becomes most evident upon metabolic reactivation. They also show that reduced *NADP-ME1* expression increases susceptibility to oxidative stress during germination, while the maintenance of low lipid peroxidation in OX lines during imbibition may contribute to their enhanced vigor and improved germination performance.

**Figure 2.**
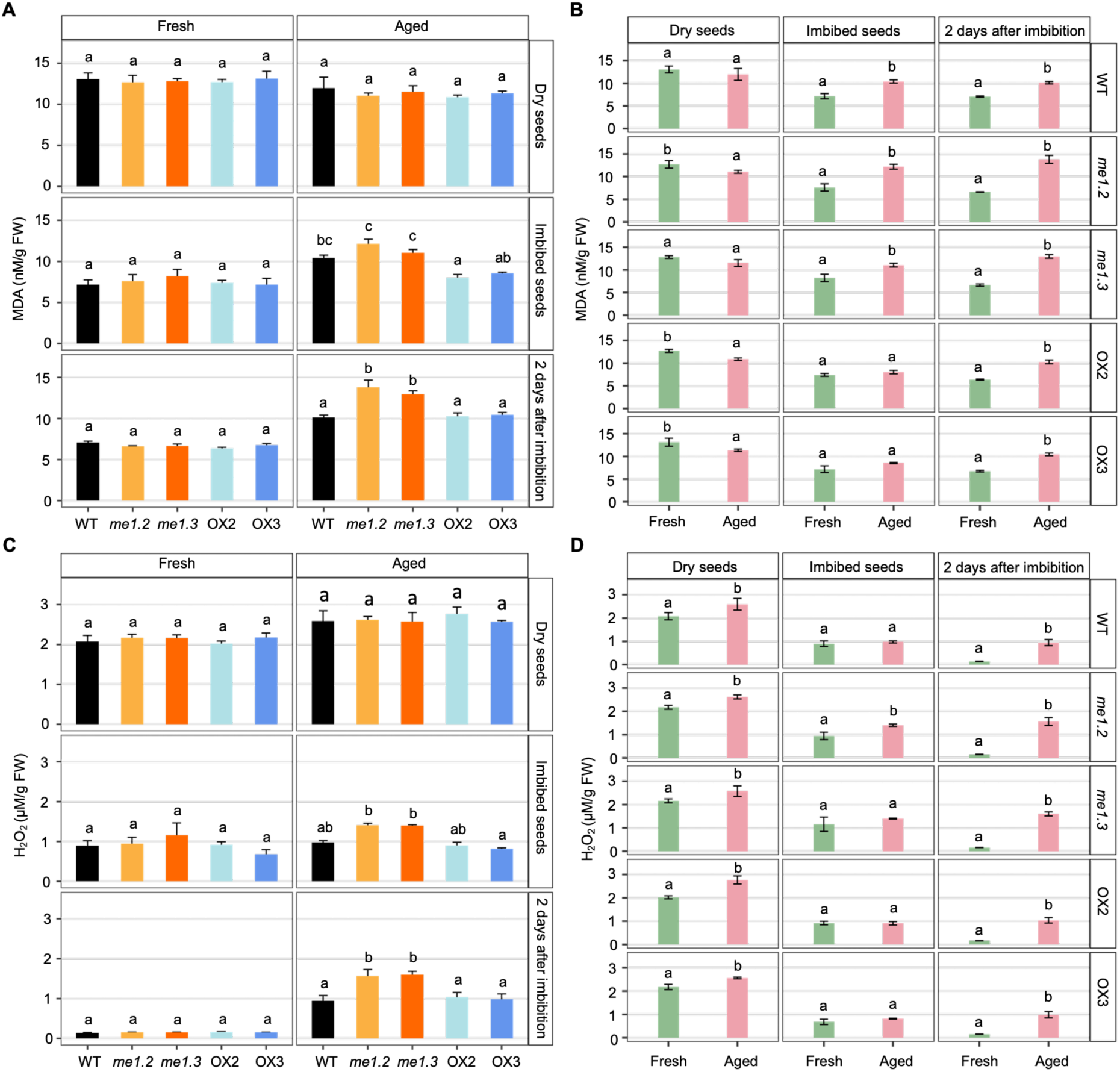
Levels of oxidative stress markers malondialdehyde and hydrogen peroxide during germination in fresh and aged seeds. The analyzed genotypes were WT, *me1.*2, *me1.3*, OX2, and OX3. Measurements were performed at three developmental stages: dry seeds, seeds imbibed for 48 h, and seeds 2 days after imbibition. **A)** Malondialdehyde (MDA) levels among genotypes in fresh and aged seeds. **B)** MDA levels in fresh versus aged seeds within each genotype. **C)** H_2_O_2_ levels among genotypes in fresh and aged seeds. **D)** H_2_O_2_ levels in fresh versus aged seeds within each genotype. Data are presented as means ± SE (n = 4). Differences were analyzed using a three-way ANOVA with genotype, treatment, and seed stage as factors. In panels A and C, pairwise comparisons among genotypes within each treatment and seed stage were performed using estimated marginal means with Sidak adjustment. In panels B and D, pairwise comparisons between treatments within each genotype and seed stage were conducted using estimated marginal means with Sidak adjustment. Different lowercase letters indicate significant differences (*p* < 0.05).

### *NADP-ME1* expression level is associated with selective changes in fatty acid composition during seed aging

Total fatty acid content did not differ significantly among genotypes (Suppl. Fig. 3A) or treatments (Suppl. Fig. 3B) in either dry or imbibed seeds, indicating that total fatty acid accumulation is unaffected by seed aging or *NADP-ME1* expression. Likewise, the relative abundance of most major and minor fatty acids remained unchanged across genotypes and treatments (Fig. 3; Suppl. Figs. 3 and 4), pointing to a generally stable fatty acid profile. The few significant differences detected were restricted to polyunsaturated fatty acids: aged dry seeds of the ko lines showed a lower content of linolenic acid (18:3) than WT and OX lines, whereas aged imbibed seeds of the OX lines showed a higher content of linoleic acid (18:2) than the WT and ko lines (Fig. 3). Because linolenic acid is particularly prone to oxidation because of its three double bonds (Reszczyńska and Hanaka, 2020), its reduction in aged dry seeds of the ko lines may reflect a greater susceptibility to aging-associated peroxidative damage. Conversely, the higher linoleic acid proportion in aged imbibed seeds of the OX lines suggests better preservation of membrane lipids during early metabolic reactivation. Together, these results indicate that NADP-ME1 does not alter total lipid reserves, but may contribute to seed vigor by preserving specific unsaturated fatty acids during aging and imbibition.

**Figure 3.**
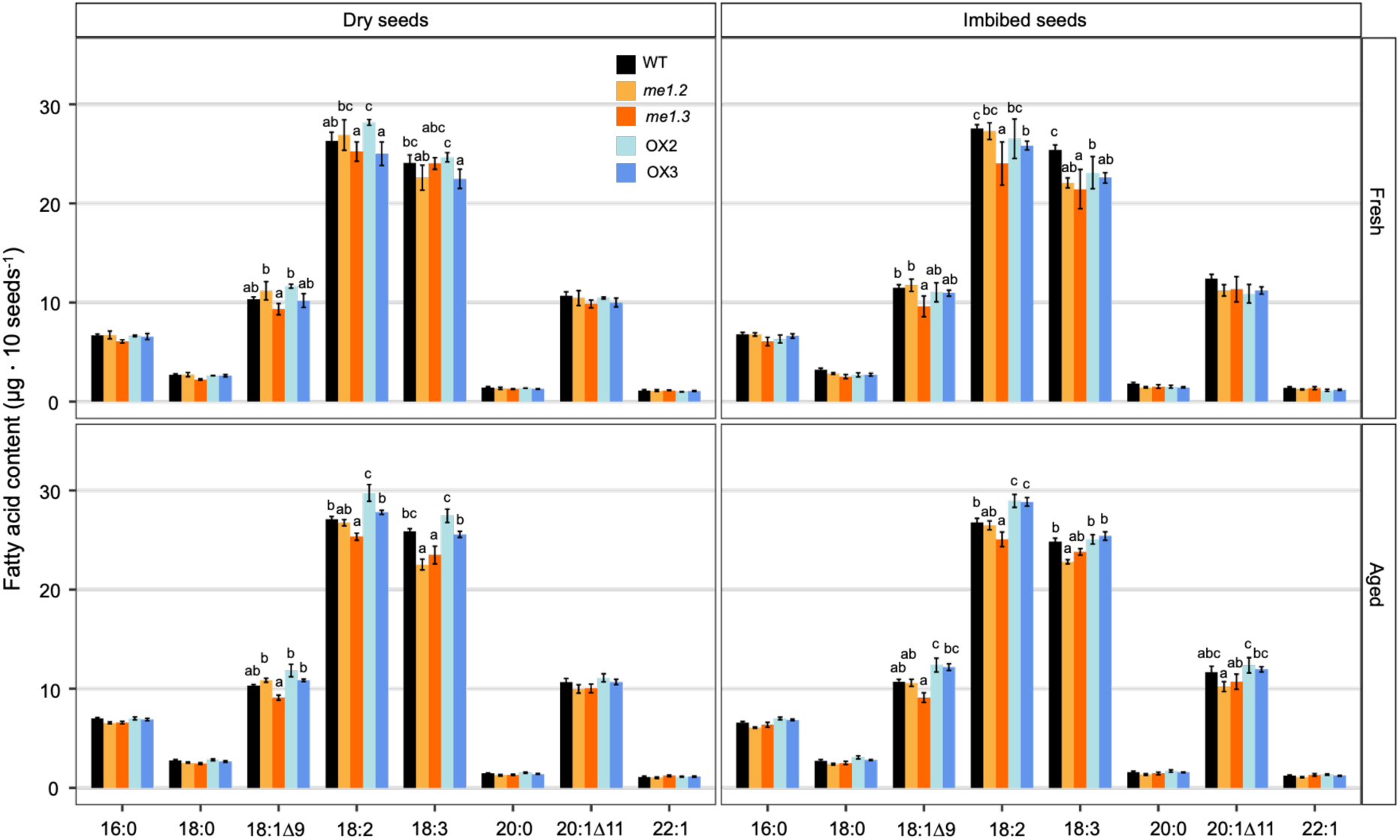
Major fatty acid composition in fresh and aged seeds. The analyzed genotypes were WT, *me1.2*, *me1.3*, OX2, and OX3. Measurements were performed at two developmental stages: dry seeds and seeds imbibed for 48 h. Contents of major fatty acids (16:0, 18:0, 18:1Δ9, 18:2, 18:3, 20:0, 20:1Δ11, and 22:1) were measured among genotypes in fresh and aged seeds. Data are presented as means ± SE (n = 5). Differences were analyzed using a three-way ANOVA with genotype, treatment, and seed stage as factors, followed by pairwise comparisons among genotypes within each treatment and seed stage using estimated marginal means with Sidak adjustment. Different lowercase letters indicate significant differences (*p* < 0.05). Letters are only shown for fatty acids that differ among genotypes; no letters are displayed when there is no significant difference.

**Figure 4.**
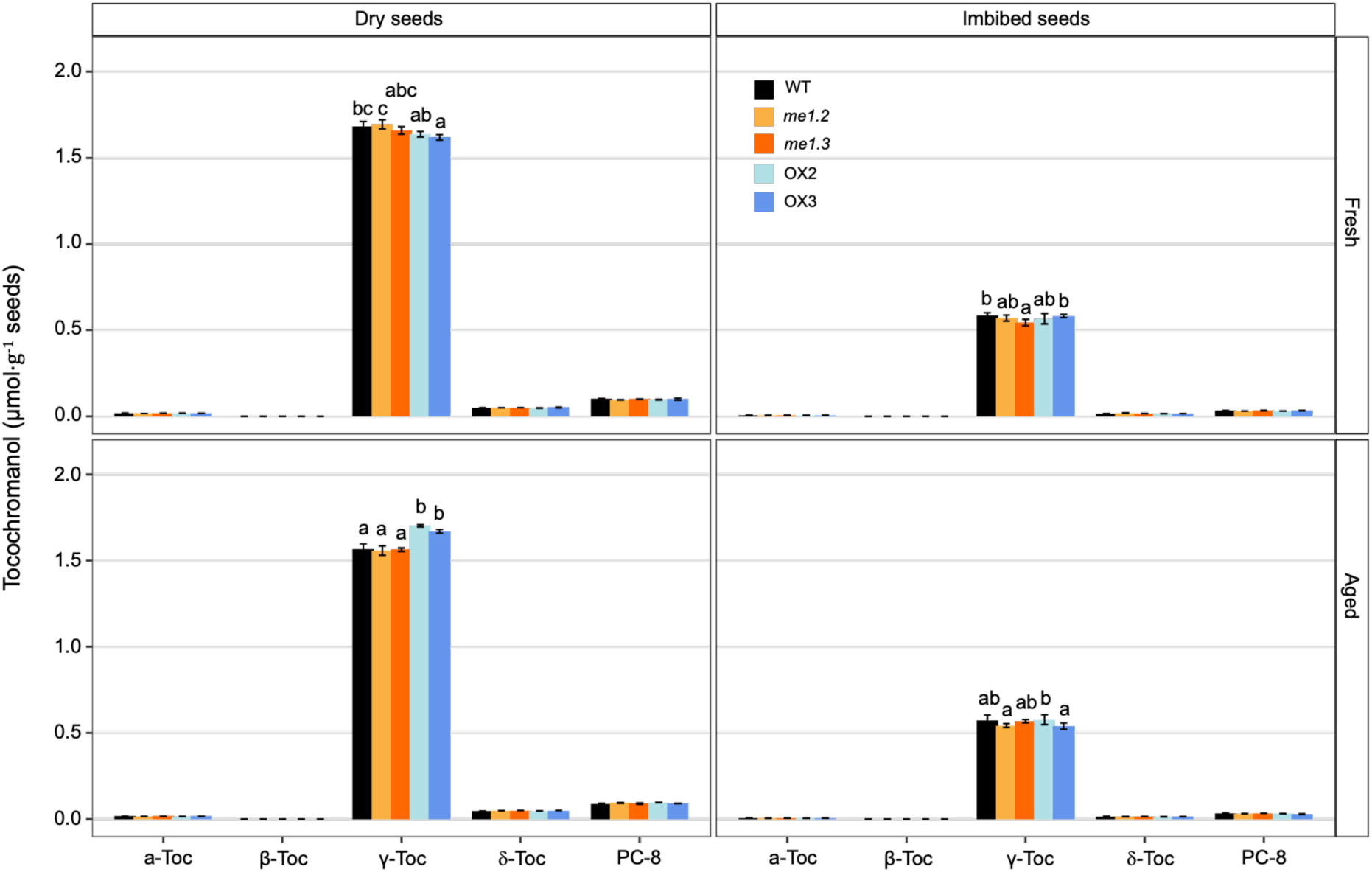
Quantification of Tocopherols in fresh and aged seeds. The analyzed genotypes were WT, *me1.2*, *me1.3*, OX2, and OX3. Measurements were performed at two developmental stages: dry seeds and seeds imbibed for 48 h. Tocopherols (α-Toc, β-Toc, γ-Toc, and δ-Toc) and plastochromanol-8 (PC-8) were measured by fluorescence HPLC among genotypes in fresh and aged seeds. Data are presented as means ± SE (n = 4). Differences were analyzed using a three-way ANOVA with genotype, treatment, and seed stage as factors, followed by pairwise comparisons among genotypes within each treatment and seed stage using estimated marginal means with Sidak adjustment. Different lowercase letters indicate significant differences (*p* < 0.05). Letters are only shown for fatty acids that differ among genotypes, and no letters are displayed when there is no significant difference.

### *NADP-ME1* overexpression promotes γ-tocopherol accumulation in aged dry seeds

We further analysed the contents of tocopherols as they represent important scavengers of lipid peroxyl radicals. We found that α-, β-, and δ-tocopherols, and plastochromanol-8 (PC-8) were present at very low levels in seeds and did not vary significantly between genotypes (Fig. 4) or treatments (Suppl. Fig. 5). γ-Tocopherol was the predominant tocopherol isoform detected in seeds and accelerated aging significantly reduced its levels in dry seeds of WT and ko lines relative to fresh seeds (Fig 4; Suppl. Fig. 5). In contrast, seeds of the OX lines showed a significant increase in γ-tocopherol under the same conditions (Suppl. Fig. 5) and accumulated significantly higher levels than other genotypes (Fig 4). Together, these results suggest that *NADP-ME1* overexpression favors the maintenance or accumulation of γ-tocopherol during dry seed aging, contributing to enhanced antioxidant protection.

### *NADP-ME1* overexpression alters genotype-dependent transcriptional states in fresh and aged conditions

To identify transcriptional features associated with *NADP-ME1* overexpression, we performed RNA-seq on fresh and aged WT and OX3 seeds harvested 2 days after imbibition. Because fresh samples had progressed further in post-germinative development than aged samples at this stage, as indicated by visible cotyledon emergence, genotype comparisons were carried out within each condition. Consistent with this developmental difference, principal component analysis (PCA) showed that the main source of variation (PC1, 90%) separated fresh and aged samples, reflecting major differences in their global transcriptional states (Fig. 5A). By contrast, PC2 explained 3% of the variance and captured a modest genotype effect within each condition.

**Figure 5.**
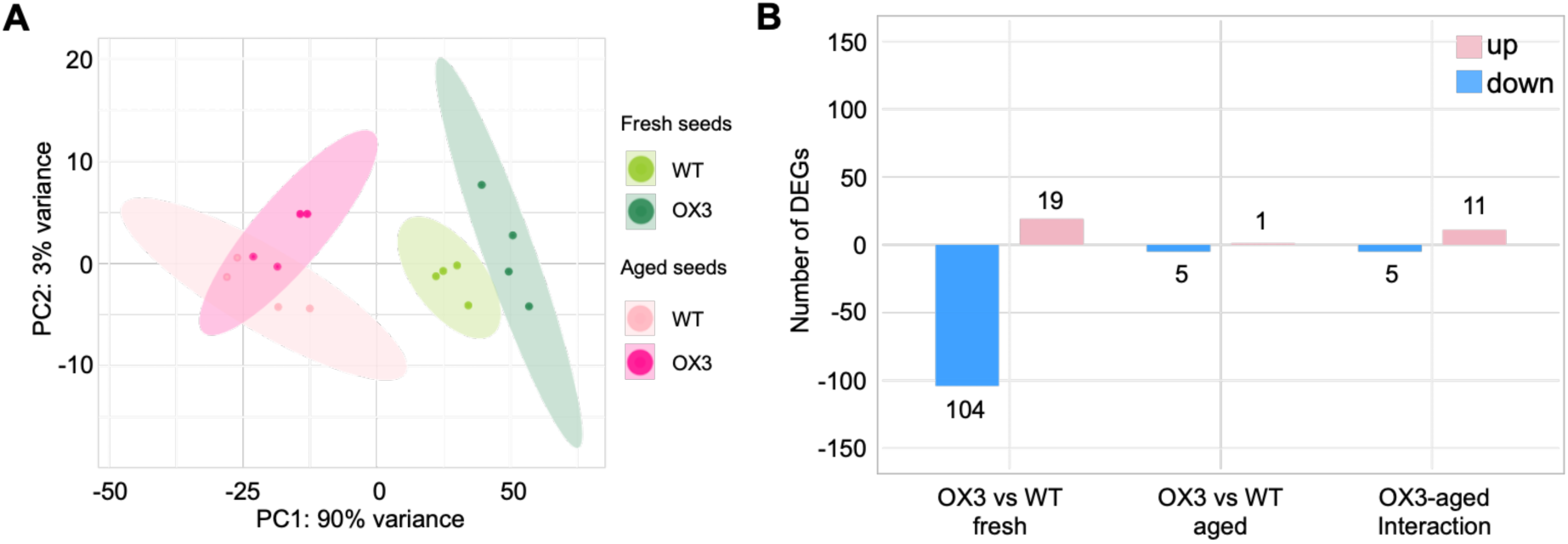
Transcriptomic responses to seed aging. **A)** Principal component analysis of transcriptomic profiles. PCA was performed on variance-stabilized normalized expression values from all samples. Each point represents one biological replicate, colored by genotype and by treatment (fresh or aged). Principal component 1 (PC1) separates samples primarily according to aging status, indicating that aging is the major source of transcriptomic variation. PC2 reflects genotype-dependent differences. **B)** Differentially significant expressed genes across interaction, state, and aging contrasts in aged and fresh seeds. Bar plots show the number of significantly up-regulated (red) and down-regulated (blue) genes for each contrast. Counts represent genes passing the significance threshold *P_adj_* < 0.05 and |log_2_FC|> 1.

Direct genotype comparisons at 2 days after imbibition identified 123 differentially expressed genes (DEGs) between OX3 and WT in fresh samples, but only 6 in aged samples (Fig. 5B). In fresh seeds, OX3-specific DEGs were marked by reduced expression of genes associated with seed maturation, storage, and dormancy, including storage proteins, oleosins, LEA proteins, and dormancy-related regulators, whereas genes involved in metabolic activity, stress adaptation, and regulatory functions were upregulated (Suppl. Data 1). Overall, this expression pattern is consistent with a more advanced post-germinative state in OX3 relative to WT, even in the absence of a visible phenotype. In aged seeds, by contrast, the limited number of DEGs suggests that the two genotypes adopt broadly similar transcriptional states despite their different germination capacities. Among this limited set of DEGs, reduced expression of stress-associated transcripts in OX3 together with increased expression of an RNA metabolism-related factor, indicating only subtle divergence in stress status and regulatory processes.

To further assess genotype-dependent transcriptional modulation across conditions, a genotype x condition interaction analysis was performed. In this multifactorial RNA-seq model, the interaction term identifies genes for which the transcriptional response to aging differs between genotypes. Within this framework, amplified genes were defined as those showing a stronger aging-induced change in OX3 than in WT, whereas attenuated genes showed a weaker response. In agreement with the PCA and DEG analyses, scatter plot analysis showed that most genes clustered near the origin, indicating that aging responses are largely conserved between genotypes. Only a small subset of genes displayed genotype-dependent modulation, including 11 amplified and 5 attenuated genes (Fig. 5B, Fig. 6A). Among the attenuated genes (Suppl. Data 1), several were related to stress-associated or metabolic functions. For example, *AT1G11350*, encoding a calmodulin-binding receptor-like kinase, and *SULTR4;2* (*AT3G12520*), involved in sulfur remobilization, showed reduced induction in OX3. Likewise, *PPO2* (A*T5G14220*), associated with tetrapyrrole metabolism, exhibited a weaker aging response in the overexpression line. *NADP-ME1* itself was also classified as attenuated, consistent with its strong induction by aging in WT and its constitutively elevated expression in OX3. By contrast, amplified genes included *SLDP1* (*AT1G65090*), associated with lipid droplet organization, *XTH5* (*AT5G13870*), involved in cell wall remodeling, and *CYP707A2* (*AT2G29090*), a key component of ABA catabolism. Additional amplified genes, including *GLXI-like8* (*AT2G28420*) and a *C2H2* zinc finger transcription factor (*AT5G48890*), further support genotype-dependent differences in detoxification and regulatory functions during aging.

**Figure 6.**
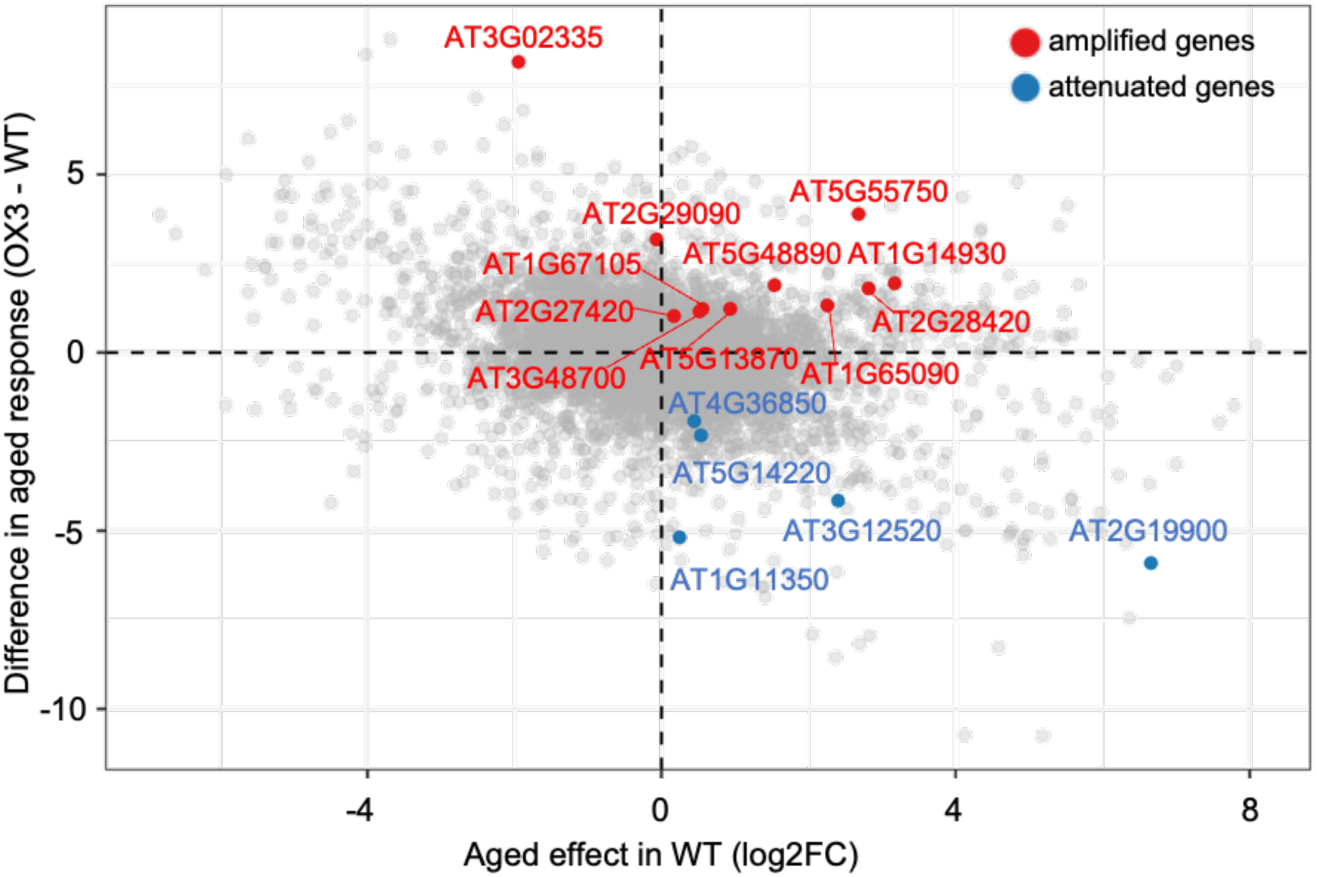
Amplified and attenuated aging-associated genes in OX3. **(A)** Scatter plot relating the transcriptional response to aging in wild-type (WT) seeds (x-axis: log_2_FC, WT aged vs fresh) to the genotype-dependent difference in aging response between OX3 and WT (y-axis: [OX3 aged − OX3 fresh] − [WT aged − WT fresh]). Each dot represents one gene. Genes with a significant genotype-aging interaction are highlighted: amplified genes (red), showing stronger aging-induced responses in OX3, and attenuated genes (blue), showing reduced responses relative to WT. Grey dots indicate genes without significant interaction effects. Dashed lines mark zero on both axes.

### Identification of NADP-ME1 interacting proteins

To gain insight into the molecular processes through which NADP-ME1 contributes to seed aging responses, we sought to identify proteins associated with it. For this purpose, we performed co-immunoprecipitation coupled with mass spectrometry (Co-IP–MS) using extracts from fresh and aged seeds of plants expressing *proME1::ME1::*YFP or *proME1::*YFP (Suppl. Fig. 5). Because *NADP-ME1* expression is higher in aged than in fresh imbibed seeds (Yazdanpanah et al., 2019), we selected the end of the imbibition phase (48 h of imbibition) to capture early NADP-ME1-associated proteins in aged seeds. At this stage, WT aged seeds sample represents an approximately equal mixture of seeds that are able to germinate and those that are not (Fig. 1C).

Among the proteins identified by MS (Suppl. Data 2), we first selected those that were strongly enriched in aged seeds in both biological replicates (log_2_ aged intensity > 2 in plants expressing *proME1::ME1::YFP* versus *proME1::YFP*), showed low abundance in fresh seeds (log_2_ fresh intensity < 1), and were localized in the cytosol (Fig. 7A). In addition, we selected candidate proteins enriched in both aged and fresh seeds (log_2_ aged intensity > 1 and log_2_ fresh intensity > 1.1) and localized in the cytosol, as these may represent constitutive NADP-ME1-associated partners and thus define a core interaction network potentially relevant to both basal seed physiology and the response to aging (Fig. 7B).

**Figure 7.**
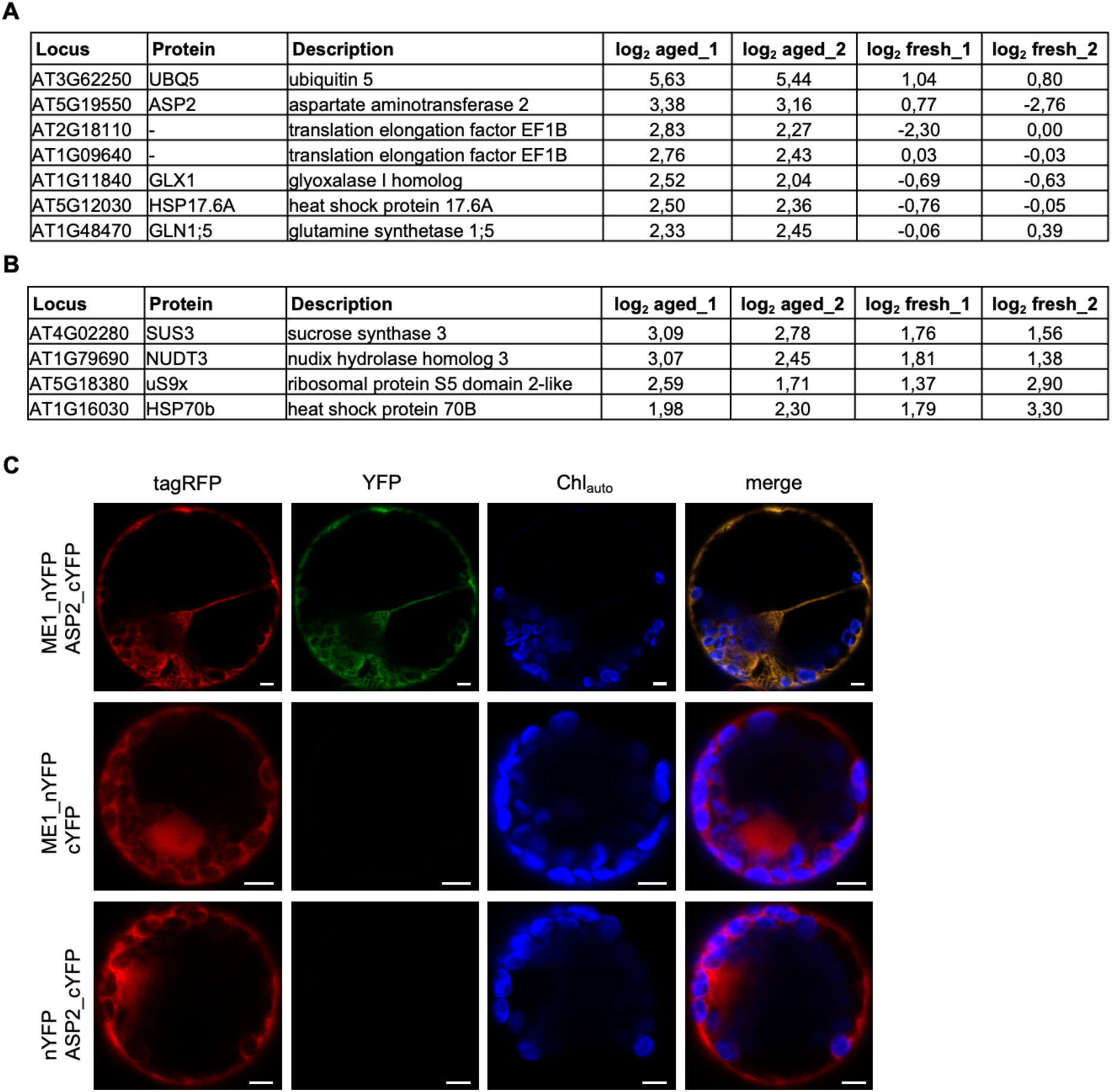
Identification and BiFC-based validation of candidate NADP-ME1 interacting proteins. **A)** Candidate proteins identified as putative NADP-ME1 interactors by co-immunoprecipitation coupled with mass spectrometry (Co-IP–MS). Cytosolic proteins were selected based on enrichment in aged seeds (log_2_ aged intensity > 2 and log_2_ fresh intensity < 1), where log_2_ aged intensity = log_2_ (*proME1*::*ME1*::YFP intensity vs *proME1*::YFP intensity) in aged seeds and log_2_ fresh intensity = log_2_ (*proME1*::*ME1*::YFP intensity vs *proME1*::YFP intensity) in fresh seeds. **B)** Additional cytosolic candidate proteins detected in both aged and fresh seeds (log_2_ aged intensity > 1 and log_2_ fresh intensity > 1.1) using the same definitions for log_2_ intensities. **C)** Bimolecular fluorescence complementation (BiFC) assays were performed by co-expressing AtNADP-ME1 fused to the N-terminal of YFP (nYFP) and candidate proteins fused to the C-terminal of YFP (cYFP) in a single 2-in-1 Gateway construct and expressing them in *Nicotiana tabacum* protoplasts. NADP-ME1 interacts with ASP2 in cytosol. Top row, Co-expression of ME1_nYFP and ASP2_cYFP showing YFP fluorescence. Second row, ME1_nYFP co-expressed with cYFP (negative control). Third row, nYFP co-expressed with ASP2_cYFP (negative control). tagRFP, red fluorescence; YFP, yellow fluorescence; Chl_auto_, chlorophyll autofluorescence; merge, overview of the protoplast. Scale bar: 5 µm.

To validate the protein-protein interactions identified by Co-IP–MS, candidate proteins were analysed by bimolecular fluorescence complementation (BiFC) in *Nicotiana benthamiana* protoplasts. Of these, only aspartate aminotransferase 2 (ASP2) interacted with NADP-ME1. Co-expression of ME1-nYFP and ASP2-cYFP yielded a strong cytosolic YFP signal, whereas no fluorescence was observed in the negative controls, including ME1-nYFP co-expressed with cYFP alone or nYFP co-expressed with ASP2-cYFP (Fig. 7C). The remaining candidates showed no detectable interaction under the conditions tested.

### Integrated network and functional analyses identify TSPO as a candidate component of the NADP-ME1 regulatory module

To uncover additional factors associated with NADP-ME1 function, we performed a co-expression network analysis using the 20 genes showing the most strongly co-expression with *NADP-ME1* (Suppl. Table 1; Fig. 8A). Gene Ontology enrichment analysis revealed a significant overrepresentation of genes associated with response to water deprivation and cold stress (Fig. 8B), further supporting a role for NADP-ME1 in stress-related seed physiology (Fu et al., 2026).

**Figure 8.**
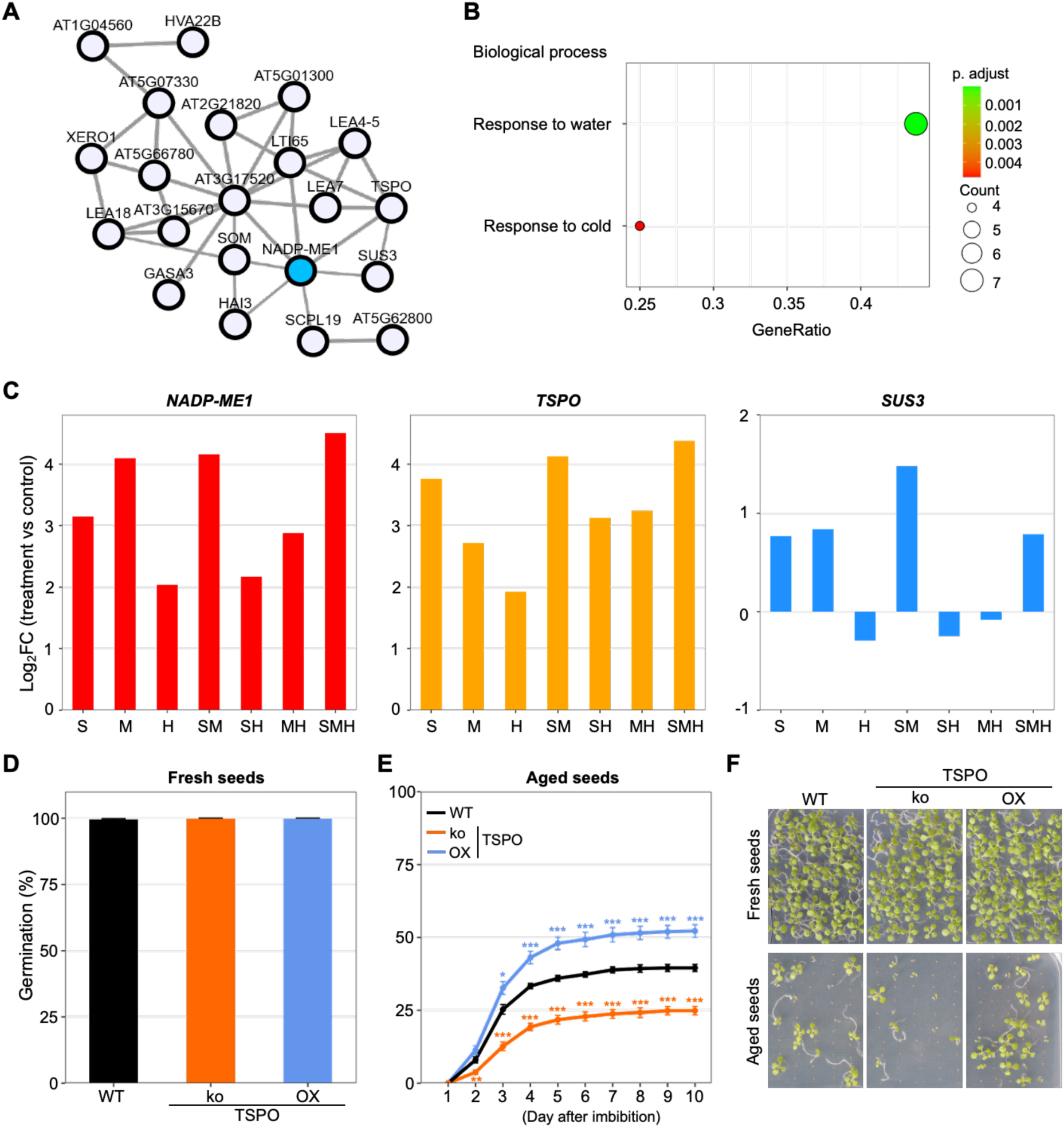
Co-expression network of *Arabidopsis NADP-ME1* and functional analysis of candidate genes. **A)** Co-expression network analysis of *NADP-ME1* performed using the ATTED-II Network Drawer tool, based on whole-genome transcriptome datasets. The network was constructed using the top 20 co-expressed genes, with *NADP-ME1* as the query gene. **B)** The Gene Ontology (GO) enrichment analysis was constructed using the top 20 co-expressed genes with *NADP-ME1* as the query gene. **C)** Relative expression levels of *NADP-ME1, TSPO* (*AT2G47770*) and *SUS3* (*AT4G02280*) under stress. Salinity (S), osmotic stress induced by mannitol (M), and heat stress (H). Expression data were obtained from Sewelam et al. (2020). Values are shown relative to control condition. **D)** Germination percentage of fresh seeds of WT, TSPO knockout mutant (ko), and TSPO overexpression line (OX) on ½ MS medium, assessed at 2 days after imbibition. **E)** Germination percentage of aged seeds from the same genotypes in D on ½ MS medium, monitored over multiple days after the imbibition phase. Germination was assessed by counting seeds with visible radicles. **F)** Representative images of fresh and aged seeds from the same genotypes in D grown on ½ MS medium for 10 days. Data are presented as means ± SE (n = 3). Asterisks indicate significant differences compared to WT (* *p* < 0.05; ** *p* < 0.01; *** *p* < 0.001; Student’s t-test).

Interestingly, SUS3, identified as a candidate NADP-ME1 interactor in the Co-IP analysis (Fig. 7B), was also identified in the co-expression network. TSPO (Tryptophan-rich sensory protein) was also detected in one Co-IP replicate (Suppl. Data 2), suggesting a potential interaction. In contrast, none of the remaining Co-IP candidates, including ASP2, were present in the co-expression network. This partial overlap suggests that while SUS3 and TSPO may belong to a broader NADP-ME1-associated functional module, the association with ASP2 may reflect a more restricted physiological context, such as seed aging. Although BiFC assays did not confirm a direct physical interaction between NADP-ME1 and either SUS3 or TSPO, their independent identification across both co-expression and Co-IP datasets strongly supports a functional relationship with NADP-ME1, potentially mediated through transient, indirect, or context-dependent associations. To further resolve these relationships, we re-examined our previous microarray dataset (Sewelam et al., 2020). *NADP-ME1* and *TSPO* showed highly similar stress-responsive expression profiles, whereas *SUS3* did not show a clear stress-related pattern (Fig. 8C). These data suggest that TSPO as part of a shared stress-responsive module with NADP-ME1, whereas the association with SUS3 may reflect a broader metabolic connection.

We next tested whether TSPO contributes functionally to the seed phenotype observed in *NADP-ME1* overexpression lines. Germination of fresh and aged seeds from *TSPO* knock-out (ko) and overexpression (OX) lines was compared with that of the WT. Under fresh seed conditions, all genotypes reached 100% germination within 2 days after imbibition (Fig. 8D). In sharp contrast, accelerated aging conditions, revealed pronounced genotype-dependent differences, as observed for NADP-ME1 (Fig. 8E and F). WT seeds showed reduced vigor, reaching only approximately 40% germination by day 7 after transfer. This impairment was further exacerbated in the ko lines, which reached only approximately 25% germination. Conversely, the OX lines displayed improved germination performance, attaining approximately 50% germination by day 7. The phenotypic similarities between TSPO and *NADP-ME1* loss- and gain-of-function lines support TSPO as a strong candidate component of the NADP-ME1 functional network.

## Discussion

### NADP-ME1 promotes seed longevity and vigorby reinforcing protection against aging-associated oxidative damage

Our findings indicate that enhanced NADP-ME1 expression helps preserve seed vigor during aging. While NADP-ME1 is dispensable for the germination of fresh seeds, its role becomes increasingly evident as seed deterioration progresses, suggesting that it acts mainly in maintenance and stress-protective processes rather than in the core germination machinery. This interpretation is supported by the reciprocal phenotypes of the genetic lines: seeds from independent overexpression lines showed higher germination efficiency during the first days after transfer and reached higher final germination after accelerated aging, whereas seeds from independent knockout lines displayed the strongest loss of vigor and longevity. Likewise, expression of a maize NADP-ME under the control of the ME1 promoter improved germination efficiency after aging, further supporting the importance of NADP-ME activity in seed protection.

The biochemical data provide a mechanistic context for this phenotype by linking NADP-ME1 to oxidative stress control. Seed aging is widely associated with increased lipid peroxidation and ROS accumulation, which compromise membrane integrity and cellular functions (Bailly et al., 1996; Kong et al., 2015; Zhang et al., 2021). Consistent with this, aging led to elevated oxidative stress, as reflected by increased MDA and H_2_O_2_ levels, particularly after metabolic reactivation in the knockout lines, while seeds of overexpression lines maintained substantially lower levels of both markers during imbibition and early germination. The absence of comparable differences in fresh seeds is especially informative, as it indicates that NADP-ME1 does not only alter the basal oxidative state, but also becomes functionally important under conditions in which oxidative damage threatens seed recovery. This is consistent with the view that the transition from quiescence to active metabolism is one of the most redox-sensitive phases of the seed life cycle (Chen et al., 2012; Rajjou et al., 2012). In this context, NADP-ME1 appears to increase the capacity of seeds to withstand the oxidative burst associated with aging and subsequent reactivation.

The increase in γ-tocopherol observed in overexpression seeds during dry aging further supports the role of tocopherols in protecting against oxidative stress. Tocopherols scavenge lipid peroxy radicals, forming tocopheroxyl radicals that can be recycled by ascorbate, thereby limiting lipid peroxidation and preserving membrane integrity (Rajjou and Debeaujon, 2008; Sattler et al., 2004). Their accumulation in overexpression lines is consistent with the reduced lipid peroxidation observed. By contrast, total fatty acid content remained unchanged across genotypes, and only minor changes in fatty acid composition were detected. Thus, the beneficial effect of NADP-ME1 cannot be explained by altered reserve accumulation, but rather by improved preservation of membrane integrity and enhanced tolerance to oxidative stress. Our results place NADP-ME1 at the center of a protective system that buffers aging-associated damage and preserves the physiological competence required for successful germination.

### Limited transcriptional reprogramming by NADP-ME1 under aging conditions

Our transcriptomic analysis conducted 2 days after imbibition reveals that the impact of *NADP-ME1* overexpression on gene expression is strongly dependent on the physiological condition and developmental progression of the samples. In fresh samples, the substantial number of DEGs and their functional distribution indicate that OX3 exhibits a transcriptional profile consistent with accelerated post-germinative progression. The coordinated downregulation of maturation- and dormancy-associated genes, together with the upregulation of metabolic and regulatory pathways, suggests that NADP-ME1 activity may promote an earlier transition from reserve-associated to growth-related programs, even in the absence of overt phenotypic differences at this stage.

In contrast, the limited transcriptional divergence observed in aged samples indicates that despite clear differences in germination performance, both genotypes converge toward a largely similar transcriptional state, suggesting that condition-specific constraints dominate gene expression and limit genotype-dependent divergence.

Consistent with this interpretation, genotype x condition interaction analysis identified only a small subset of genes with differential responses between genotypes. The predominance of conserved transcriptional patterns underscore the robustness of the condition-dependent program, while the few amplified and attenuated genes point to targeted modulation of specific pathways. In particular, altered regulation of genes associated with stress responses, sulfur and tetrapyrrole metabolism, lipid organization, cell wall remodeling, and ABA catabolism suggests that *NADP-ME1* overexpression fine-tunes metabolic and regulatory networks rather than inducing large-scale transcriptional reprogramming during aging.

Together, these findings support a model in which NADP-ME1 primarily influences early post-germinative transcriptional progression in fresh samples at 2 days after imbibition, while exerting more subtle, pathway-specific effects under aged conditions.

### NADP-ME1 integrates oxidative protection with metabolic recovery during early seed reactivation

The physiological and transcriptomic data support a model in which NADP-ME1 enhances seed vigor after aging by coordinating oxidative defense with the early reactivation of metabolism. During aging, NADP-ME1 likely contributes to antioxidant capacity, as evidenced by increased tocopherol accumulation and reduced oxidative damage. During subsequent imbibition and germination, this protection is manifested as lower ROS accumulation, better membrane preservation, and a more efficient metabolic restart. This reduction in ROS is consistent with growing evidence that NADP-ME1 operates within ABA signaling pathways to regulate ROS homeostasis, thereby linking hormonal signaling to redox control. Recent findings indicate that NADP-ME1 is required for proper ABA-dependent regulation of ROS distribution and downstream growth responses, supporting a role in actively shaping ROS dynamics rather than simply mirroring oxidative status (Fu et al., 2026). In this context, NADP-ME1-derived NADPH may fuel both ROS-generating and ROS-detoxifying systems, integrating metabolic and hormonal control of stress responses. This coordinated action may explain the faster and more successful germination of overexpression seeds after deterioration.

The identification of ASP2 as an NADP-ME1 interactor in imbibed aged seeds provides an informative mechanistic link in this model. Because this interaction was detected before visible germination, it likely reflects an early recovery process that becomes active during the first stages of seed reactivation. ASP2 catalyzes the interconversion of glutamate and oxaloacetate to generate aspartate and 2-oxoglutarate, placing it at a central hub of carbon-nitrogen metabolism (Coruzzi, 2003; Miesak and Coruzzi, 2002). This is particularly relevant because imbibed *A. thaliana* seeds are known to undergo rapid metabolic remodeling prior to visible growth, including increased levels of aspartate, threonine, serine, and 2-oxoglutarate compared with dry seeds (Arc et al., 2013; Fait et al., 2006; Nietzel et al., 2020). Moreover, aged imbibed *me1* loss-of-function seeds were previously shown to accumulate malate and amino acids (Yazdanpanah et al., 2019), consistent with a defect in coordinating carbon use and amino acid metabolism during recovery. Within this framework, the NADP-ME1-ASP2 association may represent a key node linking NADPH-generating malate catabolism with aspartate-centered carbon-nitrogen remodeling. Such a link would be highly advantageous during early seed recovery, when cells should simultaneously limit oxidative damage, re-establish metabolic flux, and prepare for resuming growth. NADP-ME1 thus emerges not simply as an antioxidant-associated enzyme, but as a broader coordinator of cellular homeostasis during the transition from aged quiescent seed to actively growing seedling.

Our data further extend this framework by identifying TSPO as a candidate component of the NADP-ME1-associated stress module. Although BiFC did not support a stable direct interaction between the two proteins, the convergence of co-expression, Co-IP, and physiological evidence supports a functional association. Similarly to *NADP-ME1*, *AtTSPO* is expressed in dry seeds and is transiently induced in vegetative tissues by osmotic stress, salt stress, or ABA treatment (Balsemão-Pires et al., 2011; Guillaumot et al., 2009). TSPO has been proposed to play a central role in the modulation of oxidative stress, largely independent of cell type or subcellular localization, at least in part through the fine regulation of tetrapyrrole metabolism (Balsemão-Pires et al., 2011; Batoko et al., 2015a; Batoko et al., 2015b). As a heme-binding protein, TSPO requires heme association for its selective degradation by autophagy. It has been proposed that the TSPO-heme interaction promotes local ROS generation, leading to TSPO oxidation, conformational change, and subsequent recognition by ATG8-related proteins (Jurkiewicz et al., 2018; Vanhee and Batoko, 2011). In this context, the similar stress-responsive expression patterns of *TSPO* and *NADP-ME1*, together with the comparable phenotypes of *TSPO* and *NADP-ME1* loss- and gain-of-function lines after accelerated aging, suggest that these proteins may operate within a shared functional framework contributing to seed vigor under deterioration stress.

While ASP2 may reflect a more specific connection to metabolic reorganization during early reactivation, TSPO appears to represent a broader stress-responsive component of this network. Together, these observations expand the functional framework of NADP-ME1 and support a model in which this enzyme contributes to the coordination of redox balance, metabolic adjustment, and stress adaptation during seed aging and germination.

### Agronomic relevance of NADP-ME1-mediated seed longevity

Beyond its mechanistic significance, the phenotype of the *NADP-ME1* overexpression lines points to agronomic relevance. Their faster and more uniform germination after aging suggests that elevated *NADP-ME1* expression could potentially improve crop establishment when seed quality has declined during storage (Finch-Savage and Bassel, 2016; Reed et al., 2022). Earlier attainment of 50% germination is particularly relevant in agricultural settings, as it is expected to promote more synchronized emergence and greater uniformity of seedling development (Ebone et al., 2020; Egli and Rucker, 2012). In practical terms, this could contribute to improved crop homogeneity, enhanced early competitiveness, and a reduced need to compensate for poor establishment through increased sowing density. The improved performance of aged overexpression seeds is also consistent with enhanced tolerance to suboptimal storage conditions, which are highly relevant for seed production, transport, and commercial handling.

Equally important, these advantages were not accompanied by broad alterations in seed physical characteristics or overall protein quality. Instead, the observed differences in protein turnover and glycation were primarily associated with the transition from the dry to the imbibed state rather than with *NADP-ME1* expression itself. From an applied perspective, this specificity is particularly attractive, as it suggests that improving vigor through NADP-ME1 does not require major trade-offs in other seed quality traits. Thus, NADP-ME1 appears to enhance seed longevity, strengthening stress resilience and recovery capacity without compromising fundamental seed properties.

### Concluding remarks

Overall, our data support a model in which NADP-ME1 acts as a central component of the seed aging response by integrating oxidative protection and metabolic recovery. Through this coordinated action, it preserves seed vigor during deterioration and enables more efficient germination after aging, making it both biologically significant and potentially valuable for crop improvement.

## Material and method

### Plant lines and growth condition

The plant lines used in this work include the ecotype Columbia-0 of *Arabidopsis thaliana* (Col-0, wild type), and the T-DNA insertion mutants of *NADP-ME1* (*AT2G19900*) previously described by Fu et al. (2026). The overexpression and knock out (Salk_135023C) lines of *TSPO* (*AT2G47770*) were kindly provided by Dr. Henri Batoko (Jurkiewicz et al., 2018). All plant lines were grown in a plant growth chamber under a 16 h light/8 h dark at 22 °C/18 °C with 120 μmol m^−2^ s^−1^ active photon flux density.

Seed longevity was assessed using an accelerated aging test as described by Sugliani et al (2009). Briefly, Aliquots of 20-25 mg seeds were placed in an open 2 mL tube and incubated in a sealed tank at 37 °C for 8 days. The tank contained a saturated KCl solution to maintain a relative humidity of approximately 83%. After incubation, the seeds were dried at room temperature for 3 days, followed by germination assay.

Seeds were sterilized with 70% ethanol for 5 min followed by 100% ethanol for 1 min, rinsed three times distilled water, and were sown on ½-strength Murashige and Skoog (MS) medium (Murashige and Skoog, 1962) solidified with 1% (w/v) agar (AGA03, formedium). Seeds were stratified for 48 h at 4 °C in the dark before being transferred to the growth condition described above. Germination was defined as seed coat rupture with radicle protrusion. For germination assay, 50 seeds per line were sown on ½ MS medium. The experiment included four technical replicates and three biological replicates.

### RNA isolation, reverse transcription, and qPCR analysis

Total RNA was isolated from roots of wild-type, T-DNA insertion mutants (*me1.1*-*me1.3*), and overexpression lines (OX1-4) using the EURX Universal RNA Purification Kit (E3598, Roboklon GmbH, Germany). For RNA-seq, total RNA was isolated using a modified SDS-LiCl method (Vennapusa et al., 2020). RNA quality and concentration were tested and measured with the NanoDrop One UV-Vis spectrophotometer (Thermo Fisher scientific) and gel electrophoresis. Genomic DNA contamination was removed using the Ambion DNA-free DNA removal kit (AM1906, Thermo Fisher Scientific). Subsequently, 750 ng of RNA was reverse transcribed into complementary DNA (cDNA) using the RevertAid Reverse Transcriptase Kit (K1691, Thermo Fisher Scientific) with an oligo(dT) primer (Suppl. Table 2), following the manufacturer’s instructions. cDNA levels were normalized to *ACTIN2* (*AT3G18780*) as an endogenous control (Czechowski et al., 2005).

For quantitative PCR (qPCR), transcript analysis of T-DNA insertion mutants and overexpression lines was performed with KAPA SYBR FAST qPCR Master Mix (KK4618, Sigma-Aldrich) using 2 µL of 1:10 diluted cDNA in a CFX96 C1000 Touch Real-Time PCR Instrument (Bio-Rad, USA), according to the manufacturer’s instructions. Primers for qPCR were designed using Primer-BLAST and SnapGene (Suppl. Table 2). Relative expression levels of *NADP-ME1* were calculated using the ΔΔCT method (Schmittgen and Livak, 2008).

### Plasmid construction and plant transformation

To produce the *pro35S::NADP-ME1* construct, the coding sequence of the *NADP-ME1* were amplified from the cDNA of wild type root using HF buffer and Phusion polymerase (F530L, Thermo scientific) with specific primers (Suppl. Table 2). The amplified fragment was sub-cloned into pCR-TOPO-Blunt (450245, Thermo scientific). The TOPO vector was used as template for further PCR-dependent fragment construction using Gibson assembly method (Gibson et al., 2009). Primers for Gibson cloning were designed to include 20-bp overlaps with the binary vector (Suppl. Table 2). The modified pGreen II vector (Fahnenstich et al., 2008) was linearized with BamHI and used in a Gibson assembly reaction with the amplified NADP-ME1 fragment. Expression of the target gene was driven by the constitutive CaMV 35S promoter. To produce the pME1:ZmnC4 construct, designed to express the maize nonC4 NADP-ME in the cytosol under the control of the AtNADP-ME1 promoter, the binary vector *pME1::ME1::YFP* (Arias et al., 2018) was modified by GenScript. Specifically, the region encompassing the *ME1::YFP* coding sequence was replaced with the cDNA of a maize NADP-ME isoform (ZmnC4, GRMZM2G122479_P01) corresponding to the mature protein, beginning at residue K43.

The resulting *pro35S::NADP-ME1* and *pME1:ZmnC4* constructs were introduced into *Agrobacterium tumefaciens* (GV3101) harboring the “helper” pSOUP plasmid via the freeze thaw method (Holsters et al., 1978). Agrobacterium tumefaciens with the target construct was transformed into Col-0 by flower dip (Clough and Bent, 1998). Transformants were selected on ½ MS medium containing the 50 mg/ml kanamycin in the case of *pro35S::NADP-ME1* or 10 mg/ml BASTA in the case of *pME1:ZmnC4*. Four homozygous T3 lines for each construction were used for further analyzed.

For bimolecular fluorescence complementation (BiFC), constructs were generated using the 2in1 Gateway cloning system (Grefen and Blatt, 2012). The *NADP-ME1* coding sequences without stop codon (-TGA) was cloned into pDONR221 P2R-P3 entry vectors (attP2 and attP3 sites) via BP Clonase™ II (11789020, Thermo Fisher Scientific). Candidate interactor was cloned without stop codon into pDONR221 P1-P4 (attP1 and attP4 sites) using the same BP reaction. Entry clones were recombined into the pBiFCt-2in1-CC destination vector using LR Clonase™ II (11791020, Thermo Fisher Scientific), generating constructs in which NADP-ME1 was fused to the N-terminal of YFP (NADP-ME1_nYFP) and interactor was fused to the C-terminal of YFP (candidate_cYFP) for subsequent BiFC assays described below.

### NADP-ME activity measurements in seed extracts

Dry fresh seeds were homogenized in mortars, and the resulting extracts were desalted through Sephadex G-50 spin columns according to Badia et al. (2015). NADP-ME activity was determined at 30 °C in a spectrophotometer (Jasco V-630) by following the increase in absorbance at 340 nm due to NADPH formation (ε340nm = 6.22 mM^-1^ cm^-1^). Assays were performed in a reaction medium containing 50 mM Tris-HCl (pH 7.5), 10 mM MgCl_2_, 0.5 mM NADP, and 10 mM malate. One unit (U) is defined as the amount of enzyme required to produce 1 μmol of NADPH per min under the specified conditions. Each sample was assayed in triplicate using three biological replicates.

### Seed Fresh Weight, Dry Weight, and Water Content Measurement

Seed fresh weight (FW) was determined by weighing 100 seeds (fresh or aged) on an analytical balance. The seeds were then dried in an oven at 65 °C for 48 h to obtain the dry weight (DW). The seed water content (%) was calculated as (FW-DW) / FW x 100.

### Protein extraction and Amadori products measurement

Proteins were extracted from 20 mg dry seeds or 50 mg of 48 h imbibed seeds. Seeds, whether fresh or aged, were ground in liquid nitrogen and homogenized in 300 µL of extraction buffer (100 mM Tris-HCl, pH 8.0; 100 mM NaCl; 0.5% (v/v) Triton X-100; 2 mM PMSF; 1% (w/v) PVP40). The homogenate was vortexed, centrifuged at 20,000 x g for 15 min at 4 °C, and the supernatant was collected. Protein concentration was determined using a modified Amido Black 10B precipitation method (Schaffner and Weissmann, 1973) with bovine serum albumin as standard for the calibration curve (Hüdig et al., 2022).

Amadori products were quantified using the nitroblue tetrazolium (NBT; N6639, Sigma-Aldrich) assay following Wettlaufer and Leopold (1991). Briefly, 0.2 mg of extracted protein was incubated with 1 mL NBT solution (0.5 mM NBT in 100 mM sodium carbonate, pH 10.3) at 40 °C. Absorbance at 550 nm was recorded after 10 and 20 min, and the increase in absorbance (ΔOD) was used to estimate Amadori product content.

### Lipid peroxidation and hydrogen peroxide quantification

Approximately 100 mg of fresh or aged seeds from each stage (dry, 48 h imbibed, and 2-day-germinated) were homogenized in 1.8 mL of 5% (w/v) trichloroacetic acid (TCA, Nr. 8789.2, Carl Roth, Germany). The homogenate was centrifuged at 1,500 x g for 10 min, and the supernatant was used for both malondialdehyde (MDA) and H_2_O_2_ quantification. Lipid peroxidation was assessed by quantifying MDA via the thiobarbituric acid (TBA; T5500, Sigma-Aldrich, USA) reaction, following Luo et al (2011) with minor modifications. Briefly, an aliquot of 0.9 mL supernatant was mixed with 0.9 mL of 0.67% (w/v) TBA, then incubated at 95 °C for 20 min, and immediately cooled on ice. Absorbance was measured at 450, 532, and 600 nm. The MDA concentration (C_MDA_, nM mL^-1^) was calculated as 6.45 x (A532 - A600) - 0.56 x A450.

H_2_O_2_ content was determined according to Velikova et al. (2000) with minor modifications. An aliquot of 0.5 mL supernatant was combined with 0.5 mL of 10 mM potassium phosphate buffer (pH 7.0) and 1 mL of 1 M potassium iodide (KI; Nr. 6750.1, Carl Roth). The mixture was incubated in darkness at room temperature for 60 min, and absorbance was measured at 390 nm. H_2_O_2_ concentration was calculated from a standard curve generated with known H_2_O_2_ solutions.

### Fatty acids quantification

Total fatty acids were measured following the method of by Browse et al (1986). Ten seeds of fresh or aged seeds at different stages (dry or 48 h imbibed) were homogenized in 2 ml safe-lock tube containing beads. Subsequently, 100 µL of internal standard (pentadecanoic acid, 15:0) and 500 µL of methanolic-HCl were added to the homogenate. The mixture was incubated at 80 °C for 30 min. After incubation, 400 µL hexane and 500 µL 0.9 % NaCl were added, followed by vortexing for 10-15 s. Samples were centrifuged at maximum speed for 1 min, and 150 µL of the upper phase was collected for gas chromatography-flame ionization detection (GC-FID) analysis. Extracts were either analyzed immediately or stored at −20 °C until analysis.

### Tocopherols quantification

Tocopherols were quantified following the method described by Vom Dorp et al (2015) with minor modifications. Approximately 20 mg of fresh or aged seeds at different stages (dry or 48 h imbibed) were ground to a fine powder. The powder was resuspended in 500 µL diethylether, and 500 ng of tocol was added as an internal standard. Phase separation was achieved by centrifugation after adding 250 µL of 0.3 M ammonium acetate in high-performance liquid chromatography (HPLC)-grade water. The (upper) organic phase was harvested, dried under nitrogen flow, then resuspended in 200 µL *n*-hexane. Tocopherol content was determined by fluorescence HPLC on a diol column, as previously described by Zbierzak et al. (2010).

### Transcriptomic analysis

Transcriptomic analysis was performed on *Arabidopsis* wild-type (WT) and NADP-ME1 overexpression (OX3) lines using fresh and aged seeds sampled at 2 days after imbibition. Accelerated aging treatment was applied prior to germination as described above. Four biological replicates were analyzed per genotype and condition. Total RNA was extracted as described above, and libraries were prepared using the Lexogen QuantSeq 3′ mRNA-Seq Library Prep Kit. Sequencing was performed on an Illumina NovaSeq 6000 platform, generating approximately 10 million single-end 100 bp reads per sample. Raw read quality was assessed using FastQC, and adapter trimming and quality filtering were performed with fastp (v0.23.2), followed by quality summarization with MultiQC. Cleaned reads were aligned to the *Arabidopsis thaliana* TAIR10 reference genome using HISAT2 (v2.2.1) with splice-aware alignment (--dta). Alignments were processed and sorted using samtools (v1.12). Gene-level read counts were generated using featureCounts (v2.0.3) with forward-stranded settings. Differential gene expression analysis was conducted using DESeq2 (v1.40.2) with a two-factor design (genotype x treatment). The following contrasts were evaluated: OX3 vs WT under aging conditions, OX3 vs WT fresh seeds, and the genotype x aging interaction effect. Genes with an adjusted *p*-value < 0.05 and |log_2_ fold change| > 1 were considered differentially expressed. Data quality and reproducibility were assessed by correlation analysis of normalized counts and principal component analysis (PCA), confirming clear separation of genotypes and treatments as well as consistency among biological replicates.

### Functional enrichment and overlap analysis

Gene set overlaps between contrasts were assessed using set operations in R. Functional enrichment analysis was performed using MapMan bin annotations. Differentially expressed genes were tested for overrepresentation in functional categories using Fisher’s exact test, followed by false discovery rate (FDR) correction. Genes expressed in the dataset (filtered count matrix) were used as background for enrichment analyses. The microarray datasets were used for the expression analysis under abiotic stress from our group previous work (Sewelam et al., 2020).

### Co-immunoprecipitation and Western Blot analysis

For co-immunoprecipitation (Co-IP) analyses, *pME1::ME1::*YFP and *pME1::*YFP seeds (Arias et al., 2018), either fresh or subjected to accelerated aging, were imbibed for 48 h and collected for protein extraction. Approximately 200 mg of imbibed seed was ground in liquid nitrogen and homogenized in 1 mL lysis buffer (50 mM Tris-HCl pH 7.5, 150 mM NaCl, 10 mM MgCl₂, 0.5 mM EDTA, 1 mM DTT, 1% Triton X-100, and 1 mM PMSF). After incubation on ice for 1 h, extracts were centrifuged for 15 min at 4 °C to remove debris. The supernatant was then incubated overnight at 4 °C with 25 µL GFP agarose beads (GFP-Trap^®^ Agarose, ChromoTek). The beads were washed with lysis buffer, and bound proteins were eluted using acidic elution buffer (200 mM glycine, pH 2.5) followed by immediate neutralization with 1 M Tris (pH 10.4). An aliquot of the eluate (10 µL) was used for western blot analysis, while the remaining samples were stored at −80 °C for subsequent mass spectrometry analysis as described below. For western blotting, proteins were mixed with 2x Laemmli buffer, heated at 95 °C for 5 min, separated by 10% SDS-PAGE, and transferred to a nitrocellulose membrane. GFP-tagged proteins were detected using anti-GFP primary antibody (1:1000, Sigma-Aldrich) and an alkaline phosphatase-conjugated secondary antibody (1:5000, Thermo Fisher Scientific). Signal detection was performed using 5-bromo-4-chloro-3-indolyl-phosphate (BCIP, No. 6368.1, Carl Roth)/NBT substrate, and images were captured using a smartphone camera.

### Proteomic mass spectrometry analysis

Eluates from GFP-Trap® Agarose were prepared for mass spectrometric analysis by in-gel digestion with trypsin essentially as described earlier (Grube et al., 2018). Briefly, eluates were shortly separated in a polyacrylamide-gel, proteins reduced with dithiothreitol, alkylated with iodoacetamide and digested overnight with 0.1 µg trypsin. Peptides were extracted from the gel, dried in a vacuum concentrator, resuspended in 17 µL 0.1 % (v/v) trifluoroacetic acid and analyzed by liquid chromatography coupled mass spectrometry as described with modifications mentions below (Prescher et al., 2021). First, peptides were trapped on a 2 cm long precolumn and then separated by a one-hour gradient on a 25 cm long C18 analytical column using an Ultimate 3000 rapid separation liquid chromatography system (Thermo Fisher Scientific). Second, peptides were injected in a Fusion Lumos (Thermo Fisher Scientific) mass spectrometer online coupled by a nano-source electrospray interface. The mass spectrometer was operated in data-dependent positive mode. Peptide and protein identification were carried out with MaxQuant version 2.5.2.0 (Max Planck Institute for Biochemistry, Germany) with standard parameters if not stated otherwise. Protein sequences from *Arabidopsis* (downloaded from “The Arabidopsis Information Resource” (TAIR): Araport11_pep from 20220914) were used for searches with standard parameters. Identified proteins were filtered: contaminants, “identified by site” proteins, reverse hits and proteins only identified with one peptide and with no quantitative values were removed.

### PEG-mediated transformation of protoplasts

PEG-mediated transformation was performed following a modified protocol from Yoo et al. (2007). *Nicotiana benthamiana* plants were grown in soil for at least 3 weeks, and young leaves were cut into strips and digested overnight in enzyme solution (0.5% Cellulase R10, 0.15% Macerozyme R10, 10 mM CaCl_2_, 0.4 M mannitol, and 5 mM MES, pH 5.6) at room temperature in the dark. The resulting mixture was filtered through a 90 µm mesh, and protoplasts were collected by low-speed (70-80 g) centrifugation and purified using flotation buffer (10 mM CaCl_2_·2H_2_O, 0.2 mM KH_2_PO_4_, 1 mM KNO_3_, 1 mM MgSO_4_·7H_2_O, 5 mM MES, 1 mM PVP-10, and 0.46 M sucrose, pH 5.6) followed by washes with W5 buffer (154 mM NaCl, 125 mM CaCl_2_·2H_2_O, 5 mM KCl, and 5 mM glucose). Purified protoplasts were resuspended in MaMg solution (0.4 M mannitol, 15 mM MgCl_2_, and 20 mM MES, pH 6.5) and mixed with 30-50 µg plasmid DNA (ME1_nYFP and candidate_cYFP 2in1 construct). Transformation was induced with 40% (w/v) PEG 1500 for 20-30 min in darkness. After treatment, protoplasts were washed with W5 buffer, centrifuged, resuspended, and incubated overnight at 22 °C in the dark before imaging. Fluorescence signals were observed using confocal laser scanning microscopy (CLSM). YFP fluorescence was excited at 488 nm, and emission was collected between 517 and 553 nm. RFP fluorescence was excited at 552 nm, and emission was collected between 579 and 633 nm. Chlorophyll autofluorescence was detected at 686-711 nm.

### Co-expression network analysis

Co-expression network analysis of NADP-ME1 performed using the ATTED-II Network Drawer tool (https://atted.jp/), based on whole-genome transcriptome datasets (Obayashi et al., 2022). The network was constructed using the top 20 co-expressed genes (Suppl. Table 1), with NADP-ME1 as the query gene. The predicted subcellular localization of these genes was obtained from the SUBA5 database (https://suba.live/). For Gene Ontology (GO) enrichment analysis, the top 20 co-expressed genes associated with NADP-ME1 were used as input, with NADP-ME1 serving as the query gene.

### Data analysis

All statistical analyses and data visualization were performed in R (version 4.4.1) using RStudio. Data were analyzed using linear models. Heteroscedasticity-robust standard errors were applied to account for potential unequal variances. Depending on the experimental design, either Student’s t-test and three-way analysis of variance (ANOVA) was used to test for significant effects. When significant effects were detected, pairwise comparisons among lines within treatment and seed stage were performed using estimated marginal means with Sidak adjustment for multiple comparisons. Data visualization was carried out using the ggplot2 package. For Figure 1 and 8, differences between lines and wild type (WT) were analyzed using Student’s t-test, while the Figures 2-4, and Supplementary Figures 1-4 were analyzed using the three-way ANOVA.

## Data availability

The data supporting the findings of this study are available from the corresponding author upon reasonable request. RNA sequencing data generated in this study have been deposited in the NCBI Sequence Read Archive (SRA) under BioProject accession PRJNA1445253 and are publicly available at https://www.ncbi.nlm.nih.gov/sra. The mass spectrometry proteomics data have been deposited to the ProteomeXchange Consortium via the PRIDE (Perez-Riverol et al., 2022) partner repository with the dataset identifier PXD076261.

## Supplementary Data

**Supplementary Data 1. Differentially expressed genes identified in RNA-seq comparisons.** Excel file containing four worksheets listing significantly differentially expressed genes (adjusted *p* value < 0.05 and |log_2_FC| > 1) for the following comparisons: NADP-ME1 overexpression line vs WT using fresh seeds, NADP-ME1 overexpression line vs WT under aged seeds conditions, and NADP-ME1 overexpression line and aging. Columns include Gene_ID (Arabidopsis locus identifier), log_2_FC, and Padj.

**Supplementary Data 2. Proteins associated with NADP-ME1 identified by co-immunoprecipitation and mass spectrometry.** Excel file containing proteins detected after co-immunoprecipitation (Co-IP) of NADP-ME1 followed by liquid chromatography-tandem mass spectrometry (LC-MS/MS). The dataset includes two independent biological replicates, Replicate 1 (samples 9-12) and Replicate 2 (samples 13-16), and lists protein identifications together with associated mass spectrometry parameters, including intensity, peptide spectrum matches, log₂ intensity, and differences in log₂ intensities.

## ACKNOWLEDGEMENTS

This work was supported by grants of the Deutsche Forschungsgemeinschaft to V.G.M. (MA2379/20-1) and by the University of Bonn through an Argelander Starter-Kit Grant to M.B.

We gratefully acknowledge the Core Facility Quantitative Lipidomics at the University of Bonn for expert technical support, in particular Dr. Georg Hölzl and Dr. Katharina Gutbrod for assistance with these measurements. Confocal imaging was performed using a Leica TCS SP8 microscope at the IZMB funded by the DFG (INST 217/939-1 FUG). We gratefully acknowledge Dr. Fatima Chigri and Dr. Susann Frank for their technical support and training in the use of the Leica SP8 confocal microscope.

The authors have no competing interests (financial/non-financial) that might be perceived to influence the interpretation of the article.

