## Supplemental Figures and Tables for "Boosting NADP-malic enzyme 1 enhances seed vigor and longevity in *Arabidopsis thaliana*"

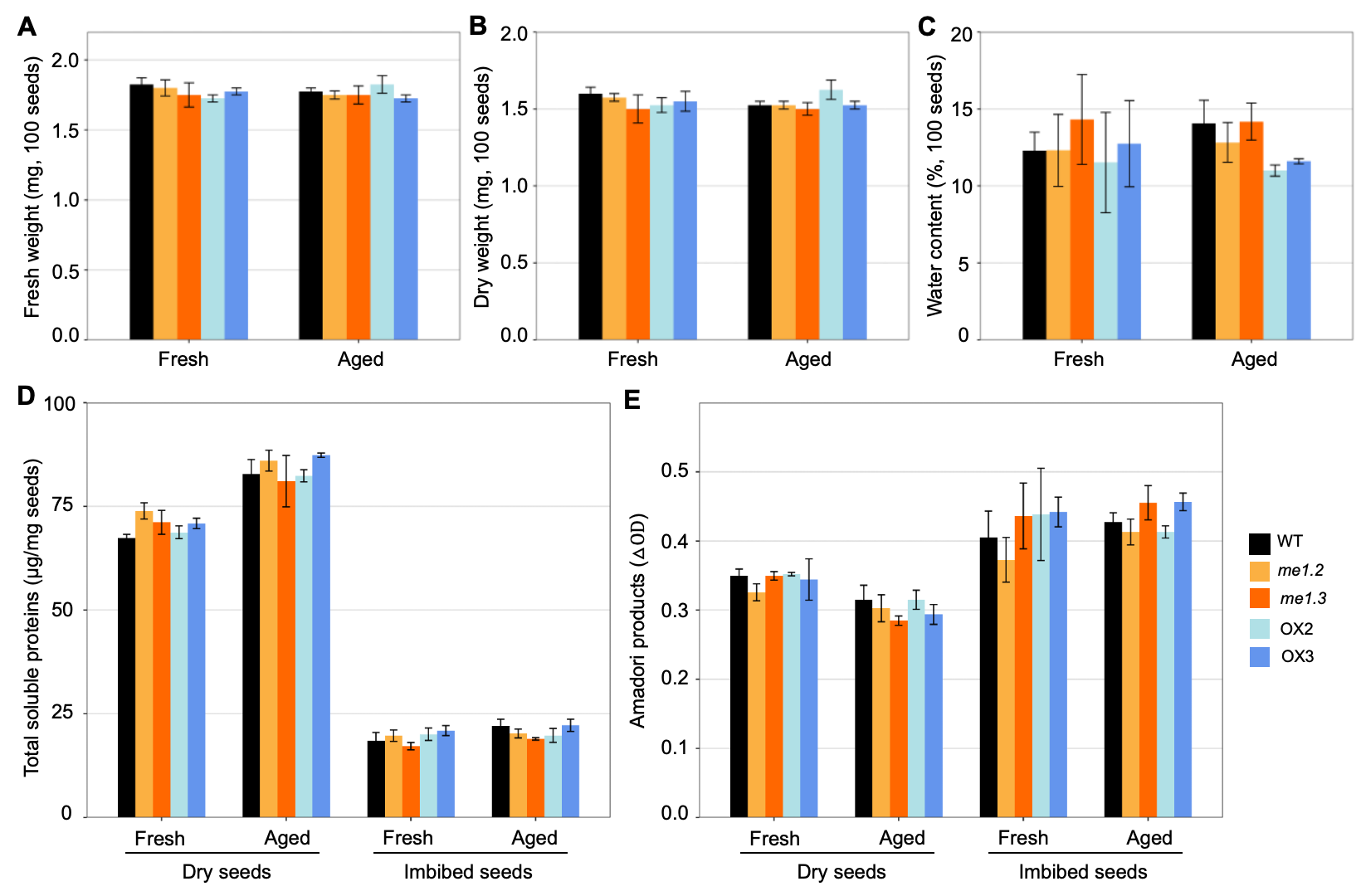

**Supplementary Figure 1. Effects of accelerated aging on seed weight, water content, protein levels, and Amadori products.** The analyzed genotypes were WT, *me1.2*, *me1.3*, OX2, and OX3. Measurements were performed at two developmental stages: dry seeds and seeds imbibed for 48 h. **A)** Fresh weight among genotypes in fresh and aged seeds. **B)** Dry weight among genotypes in fresh and aged seeds. **C)** Water content among genotypes in fresh and aged seeds. **D)** Total soluble protein content among genotypes in fresh and aged seeds. **E)** Amadori products among genotypes in fresh and aged seeds. Data are presented as means ± SE (n = 4). Differences were analyzed using a three-way ANOVA with genotype, treatment, and seed stage as factors, followed by pairwise comparisons among genotypes within each treatment and seed stage using estimated marginal means with Sidak adjustment. No significant differences were observed among the genotypes for any parameters analyzed.

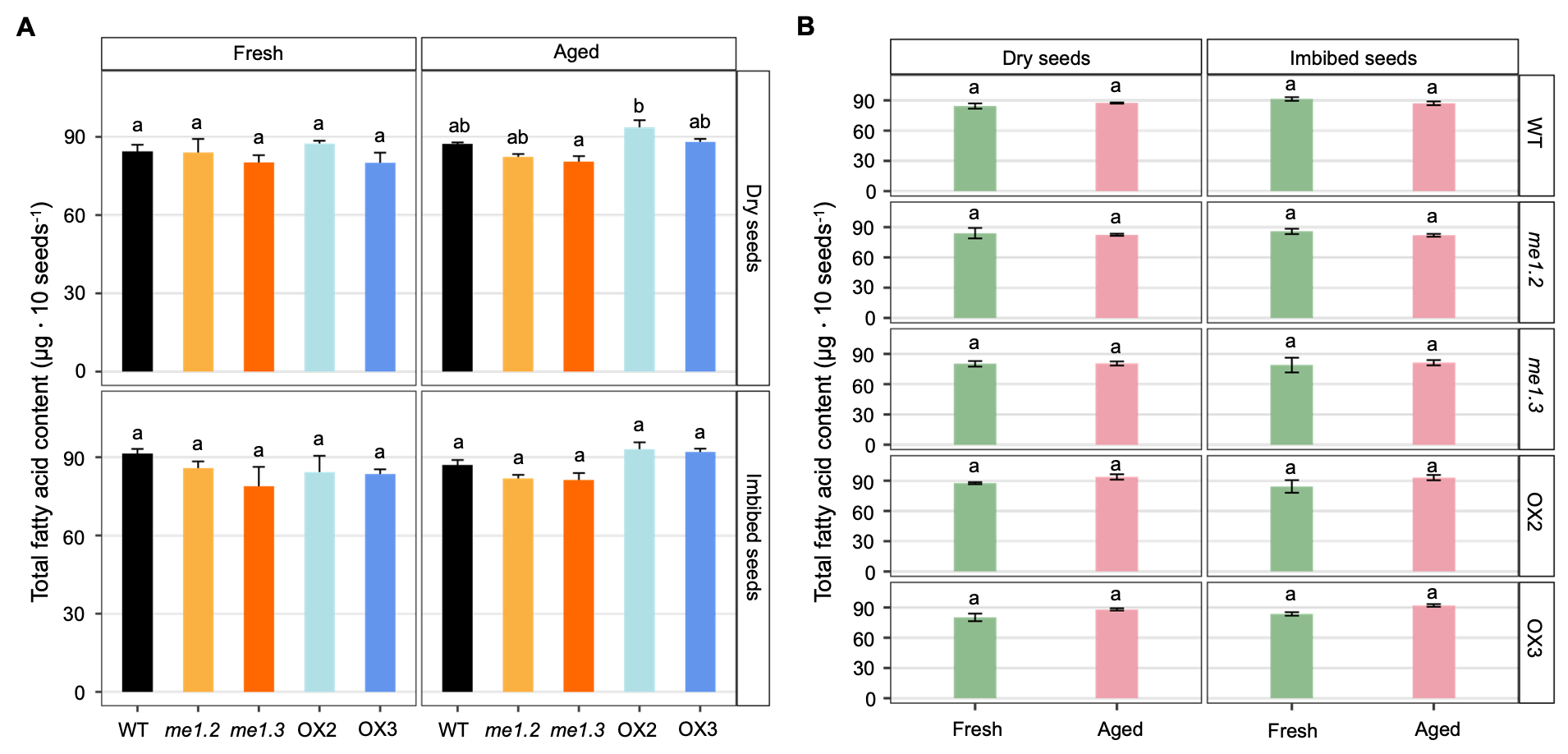

**Supplemental Figure 2. Total fatty acid content in fresh and aged seeds.** The analyzed genotypes were WT, *me1.*2, *me1.3*, OX2, and OX3. Measurements were performed at two developmental stages: dry seeds and seeds imbibed for 48 h. **A)** Comparison of total fatty acid content among genotypes in fresh and aged seeds. **B)** Comparison of total fatty acid content between fresh and aged seeds within each genotype. Data are presented as means ± SE (n = 5). Differences were analyzed using a three-way ANOVA with genotype, treatment, and seed stage as factors. In A, pairwise comparisons among genotypes within each treatment and seed stage were performed using estimated marginal means with Sidak adjustment. In B, pairwise comparisons between treatments within each genotype and seed stage were conducted using estimated marginal means with Sidak adjustment. Different lowercase letters indicate significant differences (*p* < 0.05).

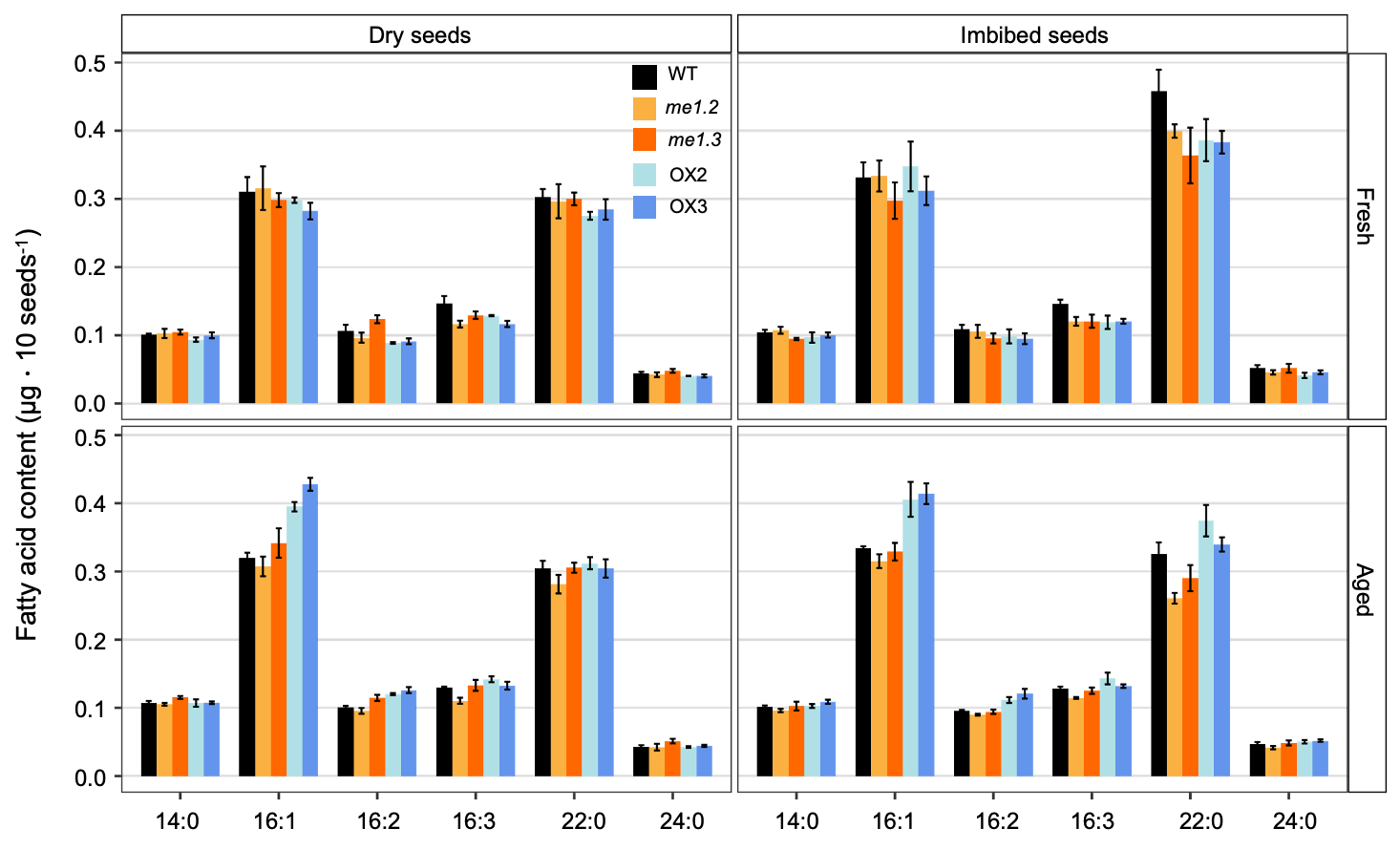

**Supplemental Figure 3. Minor fatty acid composition in fresh and aged seeds.** The analyzed genotypes were WT, *me1.*2, *me1.3*, OX2, and OX3. Measurements were performed at two developmental stages: dry seeds and seeds imbibed for 48 h. Contents of minor fatty acids (14:0, 16:1, 16:2, 16:3, 22:0, and 24:0) were measured among genotypes in fresh and aged seeds. Data are presented as means ± SE (n = 5). Differences were analyzed using a three-way ANOVA with genotype, treatment, and seed stage as factors, followed by pairwise comparisons among genotypes within each treatment and seed stage using estimated marginal means with Sidak adjustment. No letters are shown in the figure, as no significant differences were detected among genotypes for any minor fatty acids.

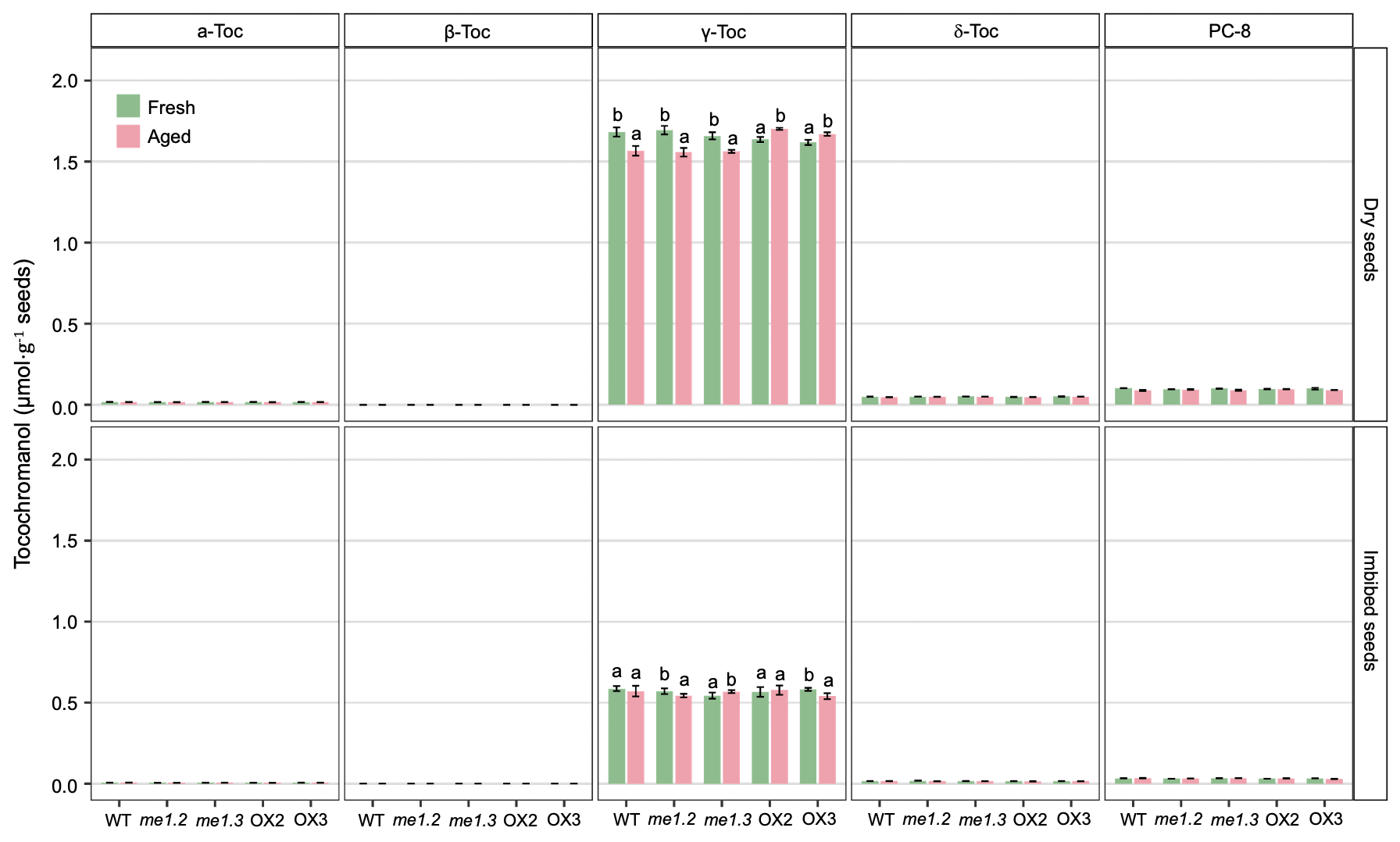

**Supplemental Figure 4. Quantification of Tocopherol in in fresh and aged seeds.** The analyzed genotypes were WT, *me1.2*, *me1.3*, OX2, and OX3. Measurements were performed at two developmental stages: dry seeds and seeds imbibed for 48 h. Tocopherols (α-Toc, β-Toc, γ-Toc, and δ-Toc) and plastochromanol-8 (PC-8) were measured by fluorescence HPLC among genotypes in fresh and aged seeds. Data are presented as means ± SE (n = 4). Differences were analyzed using a three-way ANOVA with genotype, treatment, and seed stage as factors, followed by pairwise comparisons between treatments within each genotype and seed stage using estimated marginal means with Sidak adjustment. Different lowercase letters indicate significant differences (*p* < 0.05). Letters are only shown for fatty acids that differ among genotypes, and no letters are displayed when there is no significant difference.

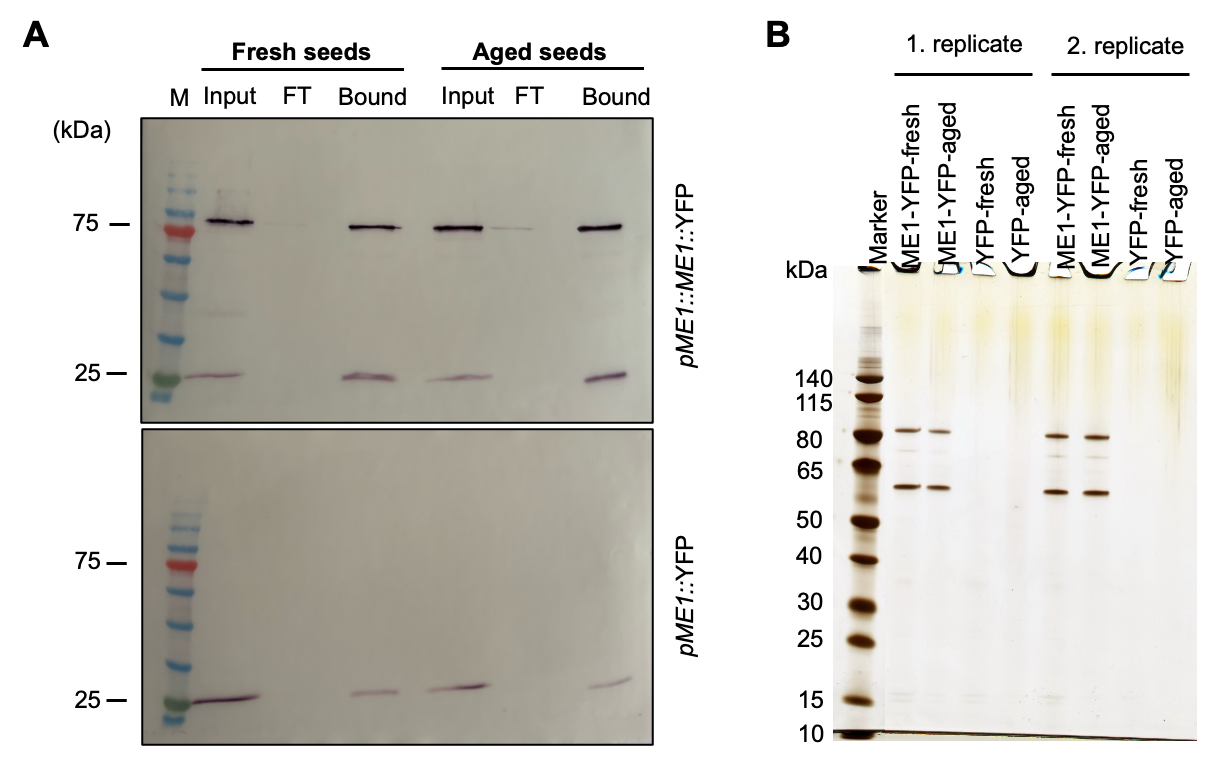

**Supplementary Figure 5. Co-immunoprecipitation (Co-IP) analysis of NADP-ME1 interacting proteins in fresh and aged seeds. A)** Protein fractions obtained from co-immunoprecipitation (Co-IP) assays using extracts in fresh and aged seeds of plants expressing *proME1*::*ME1*::YFP or *proME1*::YFP and harvested after 48 h of imbibition. Proteins were immunoprecipitated with GFP-Trap® Agarose and detected using a mouse anti-GFP (IgG1κ) primary antibody and a goat anti-mouse IgG secondary antibody. M, ROTI^®^Mark Tricolor protein marker; FT, flow-through fraction. **B)** SDS-PAGE analysis of proteins co-immunoprecipitated with ME1-YFP or YFP from the bound fractions shown in (A). One-tenth of the total volume of each bound fraction was loaded per lane. Results from two independent biological replicates are shown.

**Supplemental Table 1. List of the top 20 genes co-expressed with NADP-ME1 based on the ATTED-II unified Arabidopsis dataset (ath-u.4).**

| Gene | Other ID | Location SUBAcon | Function | ath-u.4 / NADP-ME1 | Gene ontology |
| --- | --- | --- | --- | --- | --- |
| AT3G17520 | AT3G17520 | extracellular | Late embryogenesis abundant protein (LEA) family protein | 7,8 | response to water |
| LTI65 | AT5G52300 | nucleus | CAP160 protein | 7,3 | response to water, response to cold |
| TSPO | AT2G47770 | golgi, endoplasmic reticulum | TSPO(outer membrane tryptophan-rich sensory protein)-like protein | 7,1 |  |
| SOM | AT1G03790 | nucleus | Zinc finger C-x8-C-x5-C-x3-H type family protein | 6,9 |  |
| AT5G07330 | AT5G07330 | mitochondrion | NFU1 iron-sulfur cluster protein | 6,7 |  |
| XERO1 | AT3G50980 | nucleus | dehydrin xero 1 | 6,6 | response to water, response to cold |
| HVA22B | AT5G62490 | vacuole | HVA22 homologue B | 6,5 |  |
| SCPL19 | AT5G09640 | extracellular | serine carboxypeptidase-like 19 | 6,3 |  |
| HAI3 | AT2G29380 | nucleus | highly ABA-induced PP2C protein 3 | 6,2 |  |
| SUS3 | AT4G02280 | cytosol | sucrose synthase 3 | 6,1 | response to water |
| LEA7 | AT1G52690 | nucleus | Late embryogenesis abundant protein (LEA) family protein | 6 | response to water, response to cold |
| AT5G66780 | AT5G66780 | nucleus | late embryogenesis abundant protein | 5,8 |  |
| LEA4-5 | AT5G06760 | peroxisome | Late Embryogenesis Abundant 4-5 | 5,7 | response to water, response to cold |
| GASA3 | AT4G09600 | extracellular | GAST1 protein homolog 3 | 5,6 |  |
| AT5G01300 | AT5G01300 | cytosol | PEBP (phosphatidylethanolamine-binding protein) family protein | 5,3 |  |
| AT1G04560 | AT1G04560 | vacuole | AWPM-19-like family protein | 5,3 |  |
| LEA18 | AT2G35300 | vacuole, golgi | Late embryogenesis abundant protein: group 1 protein | 5,2 | response to water |
| AT3G15670 | AT3G15670 | nucleus | Late embryogenesis abundant protein (LEA) family protein | 5,2 |  |
| AT2G21820 | AT2G21820 | nucleus | seed maturation protein | 5,1 |  |
| AT5G62800 | AT5G62800 | nucleus | Protein with RING/U-box and TRAF-like domain | 5,1 |  |

Note: Predicted subcellular localization of the encoded proteins was obtained from SUBA5.

**Supplemental Table 2. Primers used in this work**

| **purpose** | **name** | **sequence 5‘-3‘** |
| --- | --- | --- |
| RT-qPCR | ME1_Q_F1 | CCACGGCCGAAAGATATTGTC |
|  | ME1_Q_R1 | GCACACAAAGTATAGACTCTTGAG |
|  | JS-ACT2-RL-F | CTTGCACCAAGCAGCATGAA |
|  | JS-ACT2-RL-R | CCGATCCAGACACTGTACTTCCTT |
| Gibson Cloning | AT2G19900 for | ATGGAGAAAGTGACCAACTCAGAC |
|  | AT2G19900 rev | TCAACGGTAGAGACGGTATGTGG |
|  | AT2G19900_fwd-G | gacagggtacccgggATGGAGAAAGTGACCAAC |
|  | AT2G19900_rev-G | caggtcgactctagagTCAACGGTAGAGACGGTATG |
| Biomolecular fluorescence complementation (2in1 gateway cloning) | AT2G19900 for | ATGGAGAAAGTGACCAACTCAGAC |
|  | AT2G19900 rev | TCAACGGTAGAGACGGTATGTGG |
|  | UBQ5_F | GATAATCTTCAGCAGCCGTTGC |
|  | UBQ5_R | TCAAGCTTCAACTCCTTCTTTCTG |
|  | uS9x_F | ATGGCGACTCAACCAGCTAC |
|  | uS9x_R | TTAACGGTAACTCTTCTGGTAACG |
|  | TSPO_F | ATGGATTCTCAGGACATCAGATAC |
|  | TSPO_R | TCACGCGACTGCAAGCTTTAC |
|  | SUS3_F | ATGGCAAACCCTAAGCTCACTAG |
|  | SUS3_R | TCAGTCATCGGCGGTTGAAG |
|  | ASP2_F | CCCACAGCTAAAAGTTGATTCTC |
|  | ASP2_R | GGTAGATCTCAATGTCACAATCC |
|  | ME1-attB3 | GGGGACAACTTTGTATAATAAAGTTGGAATGGAGAAAGTGACCAACTCAGAC |
|  | ME1-attB2_without STOP | GGGGACCACTTTGTACAAGAAAGCTGGGTGACGGTAGAGACGGTATGTGGG |
|  | UBQ5-attB1 | GGGGACAAGTTTGTACAAAAAAGCAGGCTTAATGCAGATCTTCGTGAAA |
|  | UBQ5-attB4_without STOP | GGGGACAACTTTGTATAGAAAAGTTGGGTGAGCTTCAACTCCTTCTTTC |
|  | uS9x-attB1 | GGGGACAAGTTTGTACAAAAAAGCAGGCTTAATGGCGACTCAACCAGCT |
|  | uS9x-attB4_without STOP | GGGGACAACTTTGTATAGAAAAGTTGGGTGACGGTAACTCTTCTGGTAAC |
|  | TSPO-attB1 | GGGGACAAGTTTGTACAAAAAAGCAGGCTTAATGGATTCTCAGGACATC |
|  | TSPO-attB4_without STOP | GGGGACAACTTTGTATAGAAAAGTTGGGTGCGCGACTGCAAGCTTTAC |
|  | SUS-attB1 | GGGGACAAGTTTGTACAAAAAAGCAGGCTTAATGGCAAACCCTAAGCTC |
|  | SUS3-attB4_without STOP | GGGGACAACTTTGTATAGAAAAGTTGGGTGGTCATCGGCGGTTGAAGG |
|  | ASP2_attB1 | GGGGACAAGTTTGTACAAAAAAGCAGGCTTAATGGATTCCGTCTTCTCT |
|  | ASP2-attB4_without STOP | GGGGACAACTTTGTATAGAAAAGTTGGGTGGCCGAGGCGGGTCACTGC |
| oligodT primer | oligodT | TTTTTTTTTTTTTTTTTT |
